# An individualized fMRI protocol to assess semantic congruency effects on episodic memory in an aging multilingual population

**DOI:** 10.1101/2021.09.07.459272

**Authors:** Magali Perquin, Shivakumar Viswanathan, Michel Vaillant, Okka Risius, Laetitia Huiart, Jean-Claude Schmit, Nico J. Diederich, Gereon R. Fink, Juraj Kukolja

## Abstract

The cognitive stimulation induced by multilingualism may slow down age-related memory impairment. However, a suitable neuroscientific framework to assess the influence of multilingualism on age-related memory processes is missing. We propose an experimental paradigm that assesses the effects of semantic congruency on episodic memory using functional magnetic resonance imaging (fMRI). To this end, we modified the picture-word interference (PWI) task to be suitable for the assessment of older multilingual subjects undergoing functional magnetic resonance imaging (fMRI). In particular, stimulus materials were prepared in multiple languages (French, German, Luxembourgish, English) and closely matched in semantic properties, thus enabling participants to perform the experiment in a language of their choice. This paradigm was validated in a group (n = 62) of healthy, older participants (over 64 years) who were multilingual, all practicing three or more languages. Consistent with the engagement of semantic congruency processes, we found that the encoding and recognition of semantically related vs. unrelated picture-word pairs evoked robust differences in behavior and the neural activity of parietal-temporal networks. These effects were negligibly modulated by the language used to perform the task. Based on this validation in a multilingual population, we conclude that the proposed paradigm will allow future studies to evaluate whether multilingualism aptitude engages neural systems in a manner that protects long-term memory from aging-related decline.

## 1 Introduction

In the aging population, neurodegenerative dementias are an increasing medical and socioeconomic problem. Since effective therapies are lacking, targeting the modifiable determinants of these dementias may be the key to future strategies to prevent cognitive decline. Stimulating cognitive activities are an essential part of this toolkit (Baldivia et al., 2008, Williams et al., 2020). The increased cognitive demands of practicing two or more languages (i.e., multilingualism) have been identified as a factor that could reduce the occurrence of memory impairment and dementia (Bialystok et al., 2007, Perquin et al., 2013). Current evidence for this protective role of multilingualism is based on neuropsychological assessments. However, the neural mechanisms by which multilingualism modulates age-related episodic memory processes remain poorly understood. This gap is also methodological since prior studies have focused on the demands that multilingualism places on executive control processes rather than memory (Abutalebi et al., 2012, Dash et al., 2019, Kousaie and Phillips, 2012, Lowe et al., 2021, Bialystok and Craik, 2022, Chung-Fat-Yim et al., 2019). We sought to address this methodological shortcoming in the current study. We present a task that was specifically developed to investigate neural processes related to episodic memory in an aging multilingual population using functional magnetic resonance imaging (fMRI). This task seeks to use language-specific stimuli (i.e., words) to trigger language-independent representations (i.e., semantics) in order to assess the effect of semantic context on episodic recognition memory.

Semantic context can exert a powerful influence on how information is remembered. The congruency of scenes, objects or events can elicit an amplified neural response during encoding (Packard et al., 2017, Bar, 2004, Aminoff et al., 2013) and produce multiple behavioral effects such as an increase in the speed and accuracy of subsequent recognition while also increasing false recall (Packard et al., 2017, Crafa et al., 2017, Flegal et al., 2014). However, another critical perspective on semantic contextualization that specifically involves words is with “Stroop-like” interference effects (Starreveld and La Heij, 2017).

In the original Stroop task (Stroop, 1935), the ink color of a displayed word (e.g., **ROOM**) has to be reported (e.g., blue) while ignoring the meaning of the word. Despite this instruction, there is a relative slowing of responses when the ink color and the meaning of a word are incongruent (e.g., **RED**). This delayed response suggests a conflict between the correct response (‘blue’) and the competing response (‘red’) triggered by the automatic processing of the word’s meaning. The picture-word interference (PWI) effect (Rosinski, 1977) is particularly relevant here. This phenomenon occurs in a Stroop-like task where the object displayed in a picture must be named while ignoring a simultaneously displayed distractor word. In this task, the time to name the visually depicted object (e.g., sofa) is increased by a distractor word describing a semantically related object (e.g., chair) (La Heij, 1988, Starreveld and La Heij, 2017). Furthermore, multilingualism has been shown to modulate the PWI effect because the task involves word retrieval in one language but the simultaneous inhibition of competing words of the same meaning in other languages (Ehri and Ryan, 1980, Friesen et al., 2016). Thus, a key determinant of the PWI effect is the shared meaning (i.e., semantics) between non-language information (i.e., a picture) and language-specific information (i.e., distractor word), as well as between the meaning of words in different languages.

In the current study, we developed a paradigm to leverage the semantic contextualization processes identified by the PWI phenomenon but with some crucial modifications to be suitable for an aging, multilingual population.

In our paradigm, participants were first presented with picture-word pairs that they were required to remember (**Figure 1A**). The objects described by the picture and the word on each trial could either be semantically related or not. Unlike typical PWI paradigms, our paradigm did not require either picture naming or a selective ignoring of the displayed word. Instead, participants were instructed to explicitly judge whether there was a semantic relationship between the displayed picture and word. The effect of these semantic judgments on memory was then assessed after an extended delay (> 20 minutes) by comparing the recognition of the picture-word pairs that were judged to be related versus those that were judged to be unrelated (**Figure 1B, 1C**). We consider this adapted paradigm to be better suited for studying memory-related processes in aging populations as compared to a paradigm that required stimulus information to be selectively ignored. Indeed, recent findings have shown that, relative to younger adults, older adults memorize more irrelevant distracting information irrespective of their semantic relevance (Amer et al., 2020). Furthermore, requiring participants to judge the relatedness of the picture and the displayed word retains their semantic relationship while eliminating the interference that this semantic relationship produces when the word has to be ignored.

**Figure 1.**
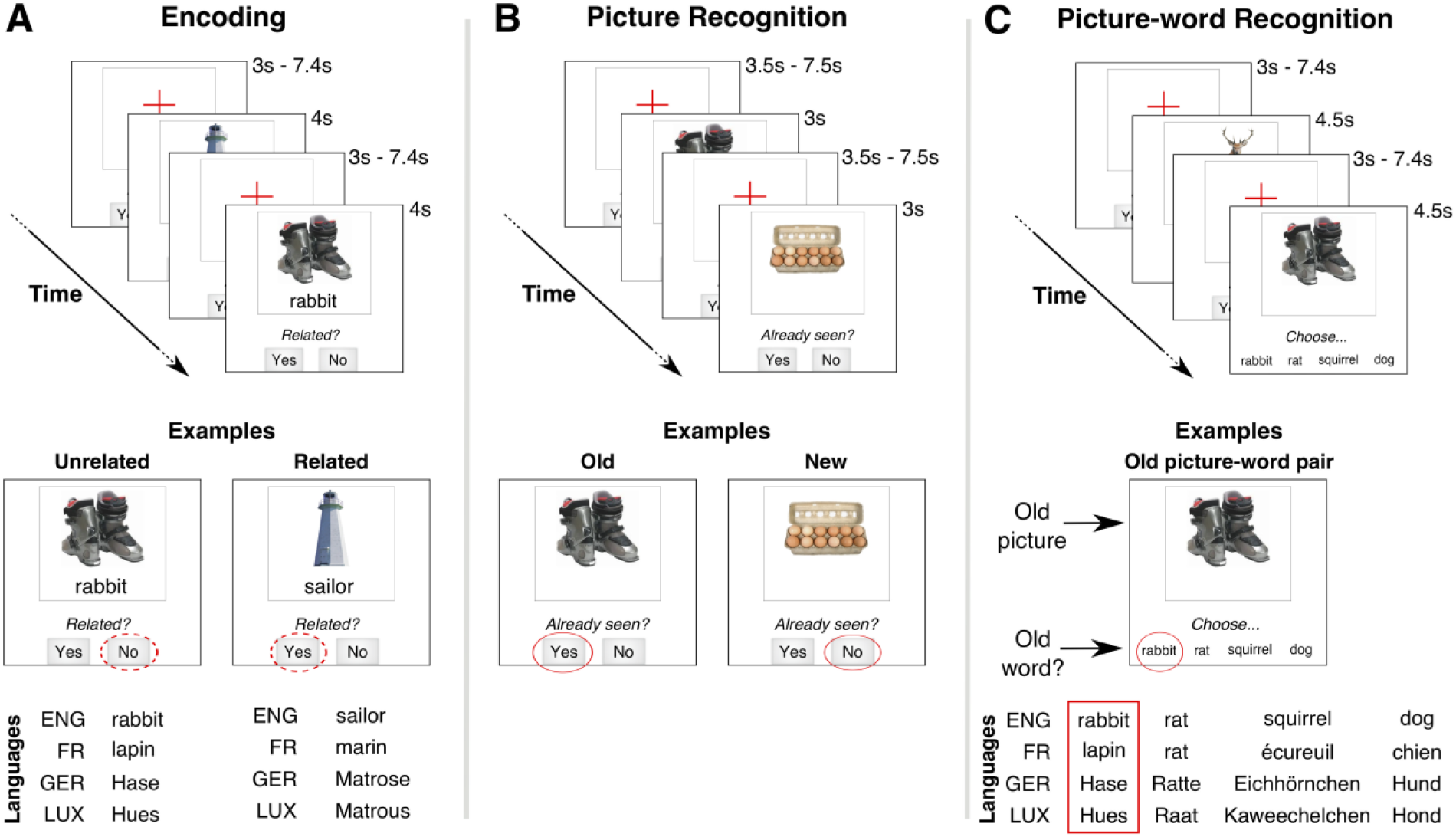
Schematic of paradigm. Trial timing and stimulus layout (not to scale) for the three task stages, with English as stimulus language. **(A) Encoding:** A picture-word pair was presented on each trial. Participants categorized as the pair as being related or not. In the examples, the ski-boots and “rabbit” could be categorized as Unrelated (“No” response) while lighthouse and “sailor” might be Related (“Yes” response). These semantic categorizations are assumed to be similar irrespective of stimulus language (ENG = English, FR = French, GER = German, LUX = Luxembourgish). **(B) Picture recognition**: A picture was presented on each trial. Participants judged whether that picture had been already seen during the Encoding task (panel **A**). In the examples, the picture of the ski-boots was previously presented (“Yes” response) but that of the egg carton was not (“No” response). **(C) Picture-word recognition:** The stimulus on each trial was a picture and four words. Participants had to choose the word that had been paired with the picture during the Encoding task (panel **A**). The three other new words were always semantically related to the target word. All the presented words were matched across stimulus languages.

Finally, a key challenge is the use of language materials with multilingual participants. One strategy to deal with a high linguistic diversity has been to use non-language materials (Abutalebi et al., 2012, Dash et al., 2019, Kousaie and Phillips, 2012). However, we reasoned that language materials would be more valuable to trigger the effects of multilingualism on cognition (Canini et al., 2016) and semantic memory. Nevertheless, an experimental protocol that uses test materials in only one language for all participants might not be suitable for a multilingual population. For example, using a single language might impose the use of a practiced but non-proficient language for some participants and also exclude participants who do not practice the testing language. In order to address these biases, we developed a multilingual stimulus library that allowed the experimental paradigm to be individually customized. Specifically, each participant was free to choose the language of the word stimuli from the four languages for which stimuli were available. To limit the scope of language-specific effects, our paradigm did not require a language-dependent response from the participant (for example, verbal picture naming). Instead, all the required responses involved choosing one of a small number of discrete alternatives with a button-press (**Figure 1**).

The value of this individualized paradigm rests on the assumption that the differential effects of semantic context (i.e., related versus unrelated) on recognition memory do not depend on the choice of stimulus language. For example, the picture of a winter hat might be displayed with the word “wool” (in English) for some participants, or “laine” (in French) for other participants, or “Wolle” (in German) for others. Judging the semantic relatedness of this picture-word pair would depend on semantic knowledge about caps, wool and their relationship (e.g., winter caps are often made of wool), which is not specific to the stimulus language. Hence, the key issue requiring empirical validation was whether a picture-word pair that is categorized as being related would be encoded and recognized differently than a pair that was categorized as being unrelated, irrespective of the stimulus language.

With this individually customized paradigm (**Figure 1**), our validation focused on addressing two central questions about the paradigm: (1) Do judgments about the semantic congruency of the stimuli during encoding (i.e., related vs unrelated) affect subsequent recognition and its associated brain activity? (2) Do the semantic congruency effects on the associated networks differ depending on the chosen stimulus language?

The current investigation was developed based on the findings of an epidemiological cohort study (Perquin et al., 2012, Perquin et al., 2013) which identified a significant association between the practice of several languages and the reduced prevalence of cognitive impairment in individuals aged 65 years and older. Therefore, our validation also focused on a similar study population of multilingual individuals who were aged 65 years and older.

## 2 Materials and Methods

### 2.1 Population selection

Sixty-two participants (mean age 70 ± 5.7 years; 36 females) completed all stages of the study protocol as part of the MEMOLINGUA study (N°REC-CESP-20141124). They were recruited using a press release. There was no financial compensation for participation. To be included in the study, participants had to be older than 64y, right-handed (self-reported), have normal or corrected-to-normal vision, have no history of psychiatric and neurological disorders, and be free of contraindications for MRI scanning.

All participants were multilingual (minimum number of practiced languages: 3). Multilingual ability was assessed with a version of the Language and Social Background Questionnaire (LSBQ) (Luk, 2008, Luk and Bialystok, 2013) that was adapted to assess multilingualism (see (Perquin et al., 2012)). This questionnaire (administered by a trained neuropsychologist) provides a detailed description of a person’s language background, self-reported proficiency in the practiced languages, and language usage patterns in different daily life contexts. Additionally, for inclusion in the main experiment, participants had to select one of the four languages for which stimuli were available as a preferred language for experimental testing (i.e., a language that they could read and respond to quickly). These four languages were French, German, Luxembourgish and English. Finally, to ensure that all participants were from a common socio-cultural context, only participants living in Luxembourg were included rather than a mixture of participants from neighboring countries such as Belgium, France, or Germany where the above languages are also practiced.

The Mini-Mental-State-Examination (MMSE) scores (Folstein, 1983) of the study population were in the normal range (mean: 28.8, SD: 1.23; min: 25, max: 30). Note that the MMSE scores were not an inclusion criterion.

The study was conducted at two locations: the Department of Population Health, Luxembourg Institute of Health (LIH), where participants were enrolled, and at the Institute of Neuroscience and Medicine (INM-3), Research Center Jülich (Germany). In one part, each participant was interviewed by a neuropsychologist at Luxembourg to obtain epidemiological information (e.g., demographics, neuropsychology, medical history, socio-cultural background, linguistic ability) following a procedure previously described in (Perquin et al., 2012). In the second part, participants were transported to (and from) the Institute for Neuroscience and Medicine, Research Centre Jülich for a single-day MRI session.

The study protocol was approved by the National Ethics Committee for Research (CNER, N°201501/03). Participants provided their signed informed consent for the entire study at LIH and for the MRI session at the Research Center Jülich.

### 2.2 Stimulus materials

Participants selected their preferred language for the stimuli to be presented during the experiment. The four available language options were Luxembourgish, French, German, and English. For each participant, the complete stimulus set across all experimental conditions consisted of 96 pictures and 256 visually depicted words in the selected language (see Supplementary Tables s1-s5).

The picture stimuli were naturalistic color photographs depicting every-day, easily nameable objects that were either natural (e.g., fruits, animals) or synthetic/artificial (e.g., tools, clothes) (Supplementary Table s5). Pictures were selected from normed databases (Brady et al., 2008, Brady et al., 2013, Konkle et al., 2010, Rossion and Pourtois, 2004).

Similar to the pictures, the words were nouns that described common natural and artificial objects. An object was not represented more than once either in the pictures or words. The word stimuli in all four languages described the same set of objects (see Supplementary Tables s1-s4). 64 (of the 96) pictures and 64 (of the 256) words were organized into unique picture-word pairs from the complete set of stimuli. These 64 picture-word pairs were further divided into two equally sized categories based on the semantic relatedness of the objects depicted by each picture-word pair. In the *Related* category, the objects described by the picture and the word pair could be easily associated with each other in a specific context in daily life. For example, a picture of a lock paired with the word “key”. In the *Unrelated* category, the objects in each picture-word pair did not share a specific contextual relationship. For example, a picture of a wine-bottle opener paired with the word “lion”. The remaining 32 (of 96) pictures and 192 (of 256) words were used as distractor stimuli in our task paradigm (see below). Specifically, each of the 64 picture-word pairs described above was associated with three distractor words. These pair-specific distractor words were always semantically related to the word of the pair and were additionally related to the picture for picture-word pairs in the *Related* (but not *Unrelated*) category.

To minimize inter-language semantic differences, we avoided picture-word relationships that might have cultural connotations specific to one language (for example, proverbs specific to French without an equivalent in German/Luxembourgish/English). A native speaker of each language evaluated the corresponding stimulus set for ambiguities.

### 2.3 Paradigm and instructions

The paradigm consisted of four tasks: (1) Encoding of picture-word pairs, (2) distractor task, (3) Picture recognition, (4) Picture-Word recognition. For clarity, only the English version of the paradigm is described below. All stimuli were displayed using Presentation® Software (Neurobehavioral Systems, Inc) on an LCD screen (size: 68.6cm (diagonal), resolution: 1200 pixels x 800 pixels, frame rate: 60 Hz). The screen was located behind the scanner and was viewed via a mirror installed on the head coil. The required responses across experimental conditions were button-presses with right hand fingers, which were recorded with an MRI-compatible LUMItouch response pad (Photon Control Inc., Burnaby, BC, Canada).

#### 2.3.1 Encoding task

On each trial of the encoding task, a visual stimulus consisting of a single picture (subtending 3.5° visual angle (v.a.)) and a single word (0.6° v.a.) below it was centrally displayed on a white screen for 4s (see **Figure 1A**). Participants had to memorize this picture-word pair. Additionally, participants judged whether the objects described by the picture and the word were related (i.e., typically associated with each other in daily life) or unrelated. This judgment was reported by pressing one of two pre-designated buttons with the right index or middle finger. Following the stimulus offset, the screen displayed a red cross during the inter-trial interval.

To continuously remind participants of the task requirements and the response mapping, the question “Related?” was displayed at the bottom of the screen (see **Figure 1A**) along with the response options “Yes” and “No”, which were positioned spatially congruent with the response buttons. During the intertrial period, this reminder was also displayed to remind participants that a response could be made even after the stimulus offset.

Special care was taken when instructing participants about the *Related/Unrelated* judgments required during the encoding task. We emphasized that the relatedness of the objects in the picture-word pair depended on whether this relationship was “typical” and not whether they could imagine a hypothetical relationship between the objects. To encourage rapid, intuitive responses, participants were told there was no correct answer for these judgments.

The stimuli for the encoding task were the 64 picture-word pairs (described above). A few additional features were incorporated to optimize the trial ordering for event-related fMRI analysis (Liu and Frank, 2004, Liu et al., 2001). Firstly, we included 32 null trials (4s duration) so that the overall task effectively had 96 trials belonging to three trial types (i.e., *Related, Unrelated*, null) with 32 trials each. The null trials were indistinguishable from the inter-trial period, and thus, for the participant, the inter-trial intervals varied over the range 3s-7.4s. Next, the 32 trials of each of the three trial types were presented in a pseudorandom order determined by a Maximum Length Sequence or m-sequence (Aguirre, 2007, Aguirre et al., 2011, Buracas and Boynton, 2002). The m-sequence ensured that all trial types were presented in a counterbalanced manner, i.e., trials of each trial type were equally likely to be preceded by trials of any trial type. The ordering of the stimuli within each trial type was randomized for each participant.

#### 2.3.2 Distractor task

To ensure that we were testing episodic recognition memory for the stimuli presented during the encoding task, an extended delay of ~24 minutes followed the encoding task. In this period, structural scans were acquired (~11 min), followed by a distractor task (~6min) and a resting state scan (~7min, see **Figure 2**). The distractor task (adapted from (Kukolja et al., 2016)) was a rapid serial visual presentation task that placed a high demand on sustained attention. This task was used to disrupt working memory processes and explicit verbal rehearsal of the stimuli encountered in the encoding task. In the task, a series of single digits from 0 to 9 were centrally presented in rapid succession (every 600ms) on a black screen, and participants pressed a button with their right index finger whenever the digit “0” was displayed. Trials were organized into 5 blocks of 48 stimuli, each separated by a 27s inter-block period. Each block contained 3 to 4 zeros.

**Figure 2.**
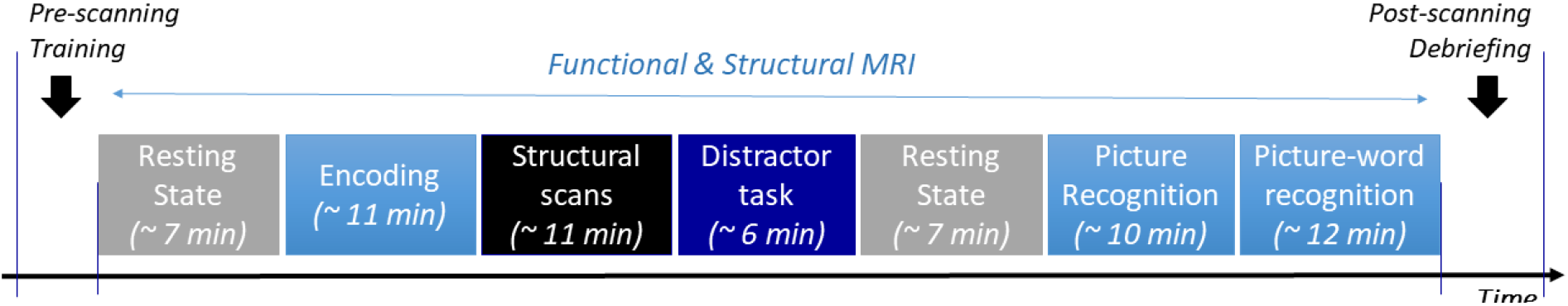
Overall procedure. Description and durations of the different consecutive stages that participants had to follow while inside the MRI scanner.

#### 2.3.3 Picture recognition task

On each trial of the picture-recognition task, a picture (3.5° v.a.) without any associated word was displayed for a 3s period (**Figure 1B**). Participants indicated whether or not they had previously seen the picture during the encoding task by pressing one of two pre-designated buttons with either the right index or middle finger. As with the encoding task, a task requirement reminder was continuously displayed at the screen’s bottom, i.e., the question “Already seen?” along with the response options “Yes” or “No”.

The stimuli for the picture recognition task were the 64 previously seen (i.e., Old) pictures from the encoding task and 32 previously unseen (i.e., New) pictures. No additional null trials were included in this task. The inter-trial interval varied randomly over the range 3.5s-7.5s. The presentation order of the 96 trials belong to the three trial-types (i.e., *Related* Old, *Unrelated* Old, New) was determined by an m-sequence, as described above for the encoding task. The ordering of the stimuli within each trial type was randomized for each participant.

#### 2.3.4 Picture-word recognition task

On each trial of the picture-word recognition task, the stimulus was a picture (3.5° v.a.) along with a horizontally displayed list of four words (0.5° v.a.) (see **Figure 1C**). Participants had to identify the word that had been previously seen with the presented picture during encoding. The correct word was always present on each trial. The identified word was indicated by the press of one of four pre-designated buttons with the right index/middle/ring/little finger. Due to a large number of items per stimulus in this task, the trial duration was increased to 4.5s. As with the encoding and picture-recognition tasks, a task demand reminder (“Choose…”) was continuously displayed at the screen’s bottom.

The stimuli for this task were the 64 picture-word pairs from the encoding task along with three semantically related distractor words (see Stimulus materials). The relative spatial position of the target word relative to the three distractors was pseudorandomized across trials, based on the Latin Square Design. As in the encoding task, we included 32 null trials (effective inter-trial interval: 3s-7.4s), and an m-sequence determined the trial ordering of the *Related*/*Unrelated*/null trial types. The ordering of the stimuli within each trial type was randomized for each participant.

#### 2.3.5 Procedure and Instructions

Each participant’s visual acuity was confirmed using a Snellen chart before the fMRI session. Prior to scanning, participants were provided with detailed written and verbal instructions on how to perform each of the different tasks. The instructions emphasized that the overall objective was to test the participant’s memory of the picture-word pairs. Furthermore, the categorization judgments were described to participants as a means to help them remember the specific pairings of pictures and words since they would be tested on their memory for these pairings. All tasks were practiced at a computer using a different stimulus set from the one used in the actual experiment to familiarize participants with the response demands and the pacing of the stimuli. This practice session was repeated until participants could comfortably perform all tasks. Participants were instructed to respond as accurately and rapidly as possible. Furthermore, we emphasized that there was no response deadline, and a response could be made even after the stimulus disappeared from view. Throughout this instruction period, we emphasized the importance of avoiding unnecessary head and body movements in the scanner. Following the scanning, participants were debriefed to assess their compliance with task instructions and fatigue levels.

### 2.4 Behavioral analysis

#### 2.4.1 Individualization of Related/Unrelated categories

Even though the picture-word pairs were designed to belong to either of two semantic categories (i.e., *Related* or *Unrelated*), participants could differ in their semantic knowledge and experience with the objects described by the stimuli. Since there was no objective accuracy criterion for these categorizations, we sought to avoid an interpretation of these subjective judgments as being “correct” or “incorrect”. Instead, all the subjective categorizations provided by an individual were treated as being “correct” for that individual. With this rationale, the picture-word pairs in the encoding task were re-categorized based on each participant’s subjective judgments about picture-word relatedness. Thus, all analyses assessing the behavior and brain activity evoked by the *Related*/*Unrelated* stimulus categories were based on each individual’s unique definition of these categories.

Since the pre-designed categories were used to optimize the trial ordering for fMRI analysis (see paradigm above), the degree of agreement between our pre-designed stimulus categorization and the individual-specific categorization was an important concern. Therefore, the degree of agreement was quantified as the proportion of trials where a participant’s judgment of picture-word relatedness (during encoding) corresponded to the pre-designed categorization of relatedness.

#### 2.4.2 Recognition accuracy and Response Times

Due to the relatively low false alarm rates (i.e., incorrect New judgments) in the picture-recognition task, we used the balanced accuracy (Brodersen, 2010) to measure overall performance rather than the classical *d-prime* measure. The balanced accuracy was equal to (H + CR)/2 where H is the hit rate (i.e., the proportion of correctly recognized Old pictures) and CR is the correct rejection rate (i.e., the proportion of new pictures that were correctly judged as being New). For all accuracy calculations, the failure to respond to a stimulus was treated as an error.

The Response Time (RT, i.e., the elapsed time from stimulus onset to the response) was only estimated for correct responses in all experimental conditions. To obtain a robust estimate of the characteristic RT for each individual in each condition, we calculated the mean RT after trimming the highest and lowest RT values (10% at each end) (Zandt, 2002, Ratcliff, 1993, Wilcox and Keselman, 2003). For completeness, the untrimmed RTs are reported in Supplementary Table s7).

#### 2.4.3 Inter-language differences

Participants were divided into separate groups depending on the language that they selected for the stimuli. To statistically assess inter-group differences in task performance, a mixed analysis of variance (ANOVA) was used to evaluate the extent to which the behavior (i.e., accuracy and RT) in each of the experimental conditions (encoding, picture-recognition, and picture-word recognition) could be independently explained by (within-subject) semantic relatedness (*Related*, *Unrelated*) and by the (between-subject) stimulus language.

Other epidemiological factors could potentially drive inter-group differences (if present). Although a large amount of epidemiological data were collected per participant, we only considered parameters of immediate relevance to the experiment itself, namely, age, gender, number of years of education, MMSE score, and the number of practiced languages (self-reported). In-depth analysis on the amount of practiced multilingualism in daily life is not part of the present study.

#### 2.4.4 Statistical analyses

Means are reported with standard deviations. For all statistical tests, normality was evaluated using the Kolmogorov-Smirnow test (when n > 50) and the Shapiro Wilk test (when n < 50). For normal distributions, we used a t-test (paired or unpaired, *t*). For non-normal distribution, non-parametric tests were used: (a) The Wilcoxon signed-rank test (*Z*) for within-subject comparisons; (b) the Wilcoxon Mann Whitney test (*U*) for two-sample comparisons. Finally, a Chi-square test (*X^2^*) was used to compare gender between participants having chosen French and those having chosen German to measure the independence of both variables in a 2-by-2 table. Behavioral results were reported in the format: *Test* (degrees of freedom) = Statistic, *p* = value. All tests were two-tailed and a p-value < 0.05 was considered statistically significant. All statistical analyses on epidemiological and behavioral data were performed with SAS System V9.4 (SAS Institute, Cary, NC, US).

### 2.5 fMRI Data

#### 2.5.1 Image acquisition and preprocessing

Functional and structural MR images were acquired on a 3T MR scanner (Siemens Tim Trio, Erlangen, Germany) with a 12-channel phased-array head coil. Functional images were measured using a T2*-weighted gradient-echo planar imaging (EPI) sequence (repetition time (TR): 2200ms, echo time (TE): 30ms, flip angle (FA): 90°, field of view (FoV): 200mm x 200mm). Each volume had 36 slices (interleaved series, thickness: 3.1mm, inter-slice gap: 0.49mm) with an in-plane resolution of 3.1mm x 3.1mm (matrix size: 64 × 64). The structural scan used a T1-weighted magnetization-prepared rapid gradient echo (MPRAGE) sequence (TR: 2250ms, TE: 3.03ms, FA: 9°, TI: 900ms, FOV: 256mm x 245mm) to obtain a high-resolution image (176 slices, matrix size: 256 × 256, inter-slice gap: 0.5 mm, voxel size: 1.0mm x 1.0mm x 1.0mm). An additional structural image was acquired using a T2-weighted fluid-attenuated inversion recovery (FLAIR) sequence, but this image was not used for preprocessing/analysis. Each task was performed on a separate scanning run. Each functional scan began with a 6TR task-free period to ensure that the MR signal reached a steady state. Since the BOLD signal evolves over multiple seconds, each functional run also ended with a 6TR task-free period to measure brain activity evoked by the final stimuli.

Image preprocessing and statistical analysis were performed with the SPM12 software (Wellcome Centre for Human Neuroimaging, London, UK) implemented for a MATLAB programming environment (MathWorks Inc., Natick, Massachusetts, USA). For preprocessing and statistical analyses, the acquired images were converted from the Siemens DICOM format to the NIFTI format using the dcm2nii utility (Li et al., 2016). Functional images (EPIs) were spatially realigned (to the first volume) and unwarped using the iterative realign/unwarp algorithm implemented in SPM12 to correct for head motion and associated magnetization artefacts (Andersson et al., 2001). The EPIs were then slice-time corrected (relative to the middle slice). The mean EPI was co-registered to the structural image. Using SPM12’s unified segmentation/normalization algorithm, the structural image was segmented to distinguish white and gray matter and then deformed to match a standard Montreal Neurological Institute (MNI) template brain image. The deformation fields estimated from this segmentation/normalization procedure were applied to all EPIs to transform them into standard MNI space (normalization) followed by resampling to a voxel size of 3mm x 3mm x 3mm (4^th^ degree B-spline interpolation). The normalized EPIs were smoothed with an isotropic 8mm full-width-at-half-maximum (FWHM) Gaussian kernel.

Following preprocessing, the fMRI and behavioral datasets of 2 (out of 62) participants were incomplete due to technical scanning difficulties that primarily affected the final picture-recognition task. Since the fMRI analysis involved inter-task comparisons, the datasets from these 2 participants were excluded from all fMRI analyses. However, the technical problems did not affect the behavioral data acquired from these 2 participants on the remaining tasks. Therefore, to maximize the use of available data, these 2 participants were included in all behavioral analyses except for the picture-word recognition task where a full set of correct trials was unavailable for response time calculations (see Supplementary Table s6 for full details of sample sizes).

#### 2.5.2 Statistical analysis

Statistical analyses were conducted within a conventional mass-univariate framework where the evoked hemodynamic response at each voxel was independently modeled using a general linear model (GLM). Each experimental condition (i.e., encoding, picture recognition, and picture-word recognition) was modeled as a separate session. Functional data from the distractor task were not analyzed as this task was included to disrupt working memory processes and explicit verbal rehearsal of the stimuli encountered in the encoding task (see above). For all conditions, the estimated first-level models shared the following properties. At every voxel, the neural response evoked by each trial was modeled as a boxcar with zero duration convolved with the canonical hemodynamic response function (HRF). The regressors of interest only modeled correct trials. Additional regressors of non-interest were included to account for incorrect trials. The 6 head-movement parameters estimated during spatial realignment (i.e., translation and rotation relative to the X, Y, Z axes) and the framewise displacement (i.e., the relative displacement of each volume relative to the previous volume) (Power et al., 2012) were included as covariates to account for head-movement effects. The BOLD time series at each voxel was high pass filtered (1/128 Hz) to remove slow trends.

We defined regressors corresponding to the *Related* and *Unrelated* trials to assess semantic congruency effects in the different tasks. For the picture-recognition task, the GLM included an additional regressor for the New trials. Contrasts were performed at the individual level (i.e., first level), and these contrast images (without any additional smoothing) were used for group (i.e., second-level) statistics.

All second-level statistics reported here were corrected for multiple comparisons at the threshold of p < 0.05 (cluster-level Family Wise Error (FWE)) with a cluster-forming threshold determined at p < 0.001 (uncorrected). We also report the minimum cluster size (in voxels) to meet this threshold for each of the contrasts as estimated in SPM12 (Nichols et al., 2016). When cluster-level correction produced overly large clusters, we use a stricter threshold of p < 0.05, voxel-wise Family Wise Error correction. For exploratory purposes, the conjunction between activity maps was assessed by the overlap in voxels that are part of statistically significant clusters in each map, i.e., the minimum-T criterion (Nichols et al., 2005).

To perform region-of-interest (ROI) analyses, each ROI was defined as a voxel cluster. An individual’s condition-specific activity at a ROI was obtained by extracting the activity estimates for that condition (i.e., beta values from the first-level analysis) from the voxels of the ROI. These voxel-wise values were then averaged together to obtain a single value that was treated as the individual’s condition-specific activity at the ROI.

Brain regions obtained from these analyses were identified using the SPM Anatomy Toolbox (Eickhoff et al., 2005), and surface visualization is shown using the Surf Ice software (https://www.nitrc.org/projects/surfice).

## 3 Results

### 3.1 Population profile and stimulus language

Forty-two participants (67.7%) chose German as their preferred stimulus language (i.e., the most comfortable language to read and answer rapidly) while 18 (29.0%) chose French and 2 (3.3%) participants chose English. None of the participants chose Luxembourgish as the stimulus language (see Supplementary s6 for summary of sample sizes).

The two largest subgroups (i.e., using German (GER) and French (FR) respectively) were similar in gender ratio, age, education, language ability, and cognitive status (MMSE scores). The mean values for these different factors are listed in **Table 1**. Importantly, there were no statistically significant differences in the population features of these two subgroups. Thus, any measured effects of stimulus language on task performance would not be attributable to the characteristics of the individuals that selected each language.

**Table 1.** Population feature differences between subgroups defined by preferred stimulus language. Only differences between subgroups using German (GER) and French (FR) are shown. (subgroup size for English (ENG) = 2, Luxembourgish (LUX) = 0). Means and standard deviations are reported for continuous variables. MMSE: Mini-Mental State Examination

|  | Subgroup<br>using<br>German<br>(GER) | Subgroup<br>using French<br>(FR) | Statistical tests |
| --- | --- | --- | --- |
| **Subgroup size** | 42 | 18 |  |
| **Gender ratio (f/m)** | 1.47 | 1.25 | $X^2(1) = 0.0816, p = 0.7751$ |
| **Age (y)** | $69 \pm 5$ | $71 \pm 6$ | $U(59) = 617.5, p = 0.2704$ |
| **Education (y)** | $14.7 \pm 4.0$ | $16.1 \pm 5.0$ | $U(59) = 590.5, p = 0.5064$ |
| **MMSE (score)** | $28.86 \pm 1.26$ | $28.83 \pm 1.25$ | $U(59) = 545.0, p = 0.9525$ |
| **Practiced**<br>**languages:** |  |  |  |
| **Lifelong (n)** | $5.3 \pm 1.3$ | $5.5 \pm 1.4$ | $U(59) = 527.5, p = 0.6539$ |
| **Current (n)** | $4.1 \pm 1.1$ | $4.5 \pm 1.6$ | $U(59) = 491.5, p = 0.4027$ |

### 3.2 Semantic congruency effects during picture-word encoding

We used the data from all participants (irrespective of stimulus language) to evaluate the main effect of semantic relatedness judgments on memory performance.

Participants’ subjective categorization of picture-word pairs as either *Related* or *Unrelated* showed a high degree of agreement to the pre-designed stimulus categories (94.5 ± 3.7%). This high degree of the agreement confirmed that participants had correctly interpreted our task instructions. Importantly, it also confirmed that the relatedness judgments, although subjective, were not arbitrary but highly consistent between participants and our pre-designed stimulus categories.

The mean response time (RT) for *Related* judgments (1749 ± 452 ms) was shorter than for *Unrelated* judgments (1877 ± 431 ms, *Z* [61] = 657.5, *p* < 0.0001) (**Figure 3A**). The contrast *Related* > *Unrelated* revealed higher activation, especially over the left hemisphere (**Figure 3B**, **Table 2**), including the left middle temporal gyrus (MTG), left posterior cingulate cortex (PCC), left anterior cingulate cortex (ACC), left inferior frontal gyrus (IFG), and bilaterally over the angular gyrus. Inversely, the contrast *Unrelated* > *Related* did not reveal any statistically significant activity difference across the brain.

**Figure 3.**
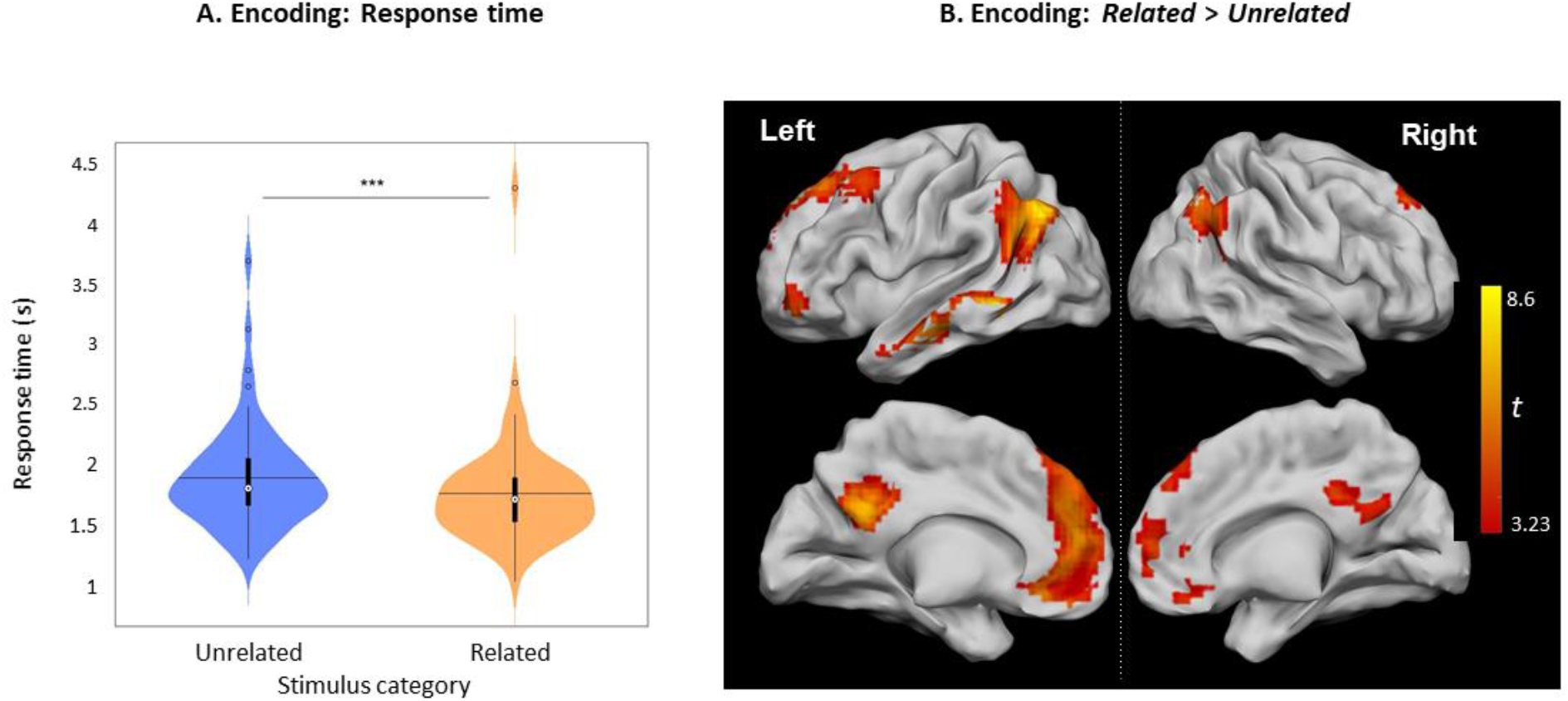
Related vs. Unrelated during encoding. **A.** Violin plots of RTs for *Related* and *Unrelated* judgments during encoding (***: p < 0.001), horizontal lines represent the mean, thick vertical black bars are a box plot, the center white circle represents the median. **B.** Clusters with increased activity during *Related* versus *Unrelated* judgments (p < 0.05, FWE-cluster corrected, cluster-forming threshold = 80 voxels (at p < 0.001 (uncorr.)).

**Table 2.** *Related* vs. *Unrelated* during encoding: Peak coordinates of significant clusters. Clusters identified at threshold of p < 0.05, FWE-cluster-corrected, cluster-forming threshold = 80 voxels (p < 0.001, uncorr.). Peak locations are in MNI coordinates. Hemisphere (Hem.) reported as L/R (Left/Right). ACC: anterior cingulate cortex, PCC: posterior cingulate cortex, IFG: inferior frontal gyrus, MTG: Middle Temporal Gyrus

| Contrast | Anatomical Region | Hem. | Coordinates |  |  | *t*-value | *p*-value | Cluster size |
| --- | --- | --- | --- | --- | --- | --- | --- | --- |
|  |  |  | x | y | z |  |  |  |
| *Rel* > *Unrel* | Angular Gyrus | L | -48 | -67 | 35 | 8.60 | 0.000 | 516 |
|  | MTG | L | -60 | -46 | -7 | 8.12 | 0.000 | 281 |
|  | PCC | L | -3 | -52 | 26 | 7.81 | 0.000 | 464 |
|  | ACC | L | -6 | 47 | 17 | 6.56 | 0.000 | 1460 |
|  | Angular Gyrus | R | 48 | -64 | 41 | 6.00 | 0.000 | 241 |
|  | IFG (p. Orbitalis) | L | -42 | 47 | -10 | 5.35 | 0.031 | 80 |
| *Unrel* > *Rel* | Nothing survived after correction |  |  |  |  |  |  |  |

### 3.3 Semantic effects on picture-recognition

Accuracy for differentiating Old and New pictures was high (89.8 ± 7.5%) and was significantly greater than random chance (50%, *U* [61] = 5797, *p* < 0.0001). Importantly, the mean accuracy in recognizing Old pictures that were part of *Related* picture-word pairs during encoding (92.5 ± 7.7%) was greater than for Old *Unrelated* pictures (82.9 ± 12.8%, *t* [61] = 7.24, *p* < 0.0001, **Figure 4A**). Furthermore, the mean RT to recognize Old *Related* pictures (1082 ± 158ms) was shorter than for Old *Unrelated* pictures (1247 ± 236ms, *Z* [61] = 905.5, *p* < 0.0001, **Figure 4B**).

**Figure 4.**
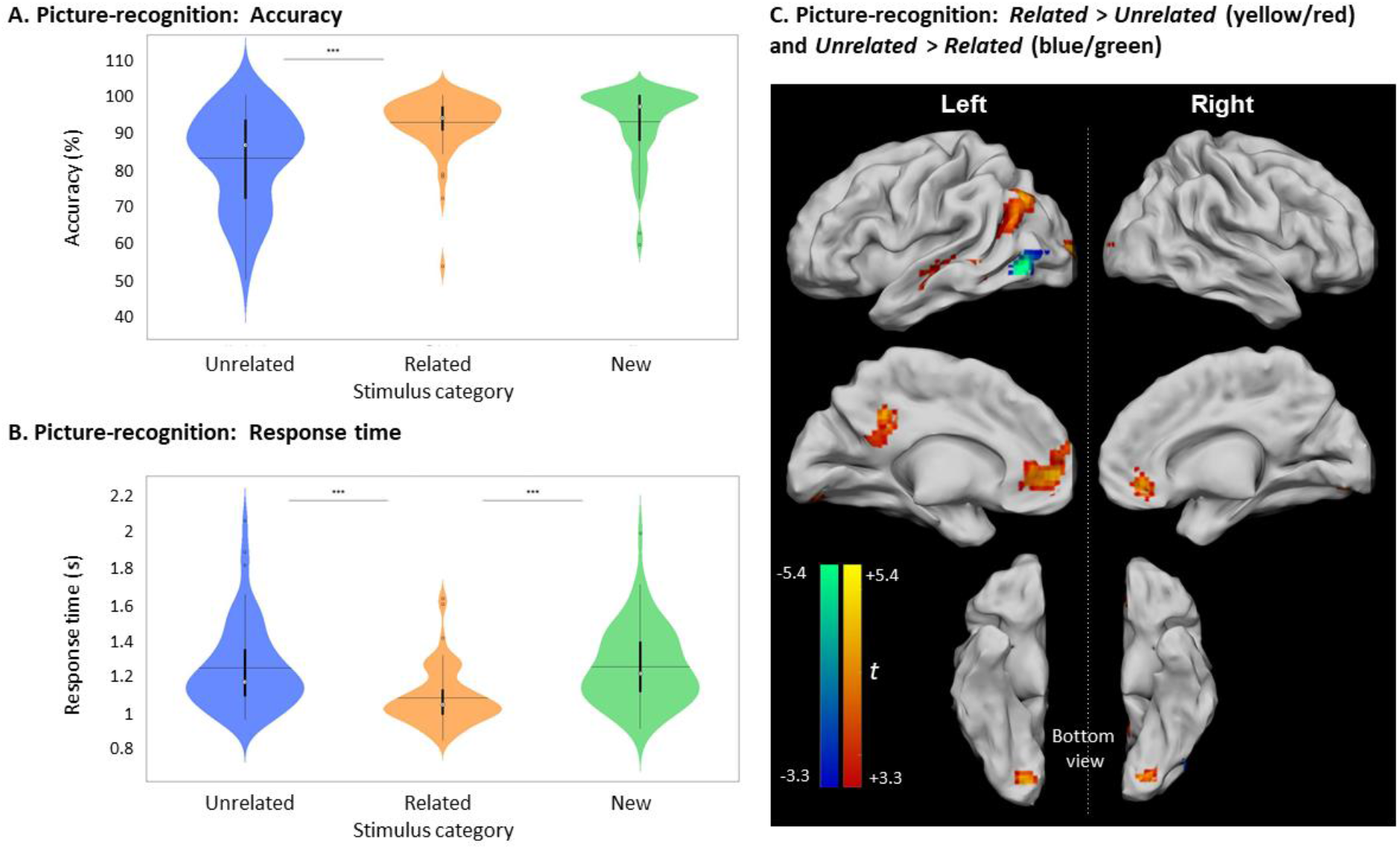
Picture recognition based on relatedness during encoding. **A.** Violin plots of recognition accuracy for New stimuli and Old stimuli, depending on their relatedness during encoding (i.e., *Unrelated* or *Related*).. [***: p < 0.001, horizontal black line indicates the mean, thick vertical black bars are a box plot, center white circles represent the median]. **B.** Violin plots of Response Times for correct New and correct Old judgments depending on their relatedness during encoding [notations as in panel **A**]. **C.** Clusters with significant activity during recognition for *Related* > *Unrelated* (red scale) and for *Unrelated > Related* (blue scale), p < 0.05, FWE-cluster corrected, cluster-forming threshold = 84 voxels (at p < 0.001, uncorr).

The accuracy in recognizing New pictures (92.7 ± 9.6%) was greater than for Old *Unrelated* pictures (*t* [61] = 4.72, *p* < 0.0001) but was not significantly different from the accuracy for Old *Related* pictures (*Z* [61] = −109.5, *p* ≥ 0.3505). However, the mean RT to recognize Old *Related* pictures was shorter than for New judgments (1253.5 ± 210.8 ms, *t* [61] = 6.17, *p* < 0.0001), while the RTs for New judgments and Old *Unrelated* pictures were not significantly different (*t* [61] = 0.20, *p* = 0.8450).

Brain activity elicited by recognizing Old *Unrelated* pictures was higher than for the Old *Related* pictures only in the left inferior occipital gyrus (**Figure 4C**, **Table 3**). However, consistent with the activity differences at encoding, the recognition of Old *Related* pictures evoked more significant activity than for Old *Unrelated* pictures over several regions, including the left MTG, the left PCC, the left ACC, the left middle occipital gyrus, and the right lingual gyrus (**Figure 4C**, **Table 3**).

**Table 3.** Picture recognition: Peak coordinates of significant clusters for all contrasts. Clusters for *Rel vs Unrel* were identified at threshold of p < 0.05, FWE-cluster-corrected cluster-forming threshold = 84 voxels (at p < 0.001, uncorr). For the *Old* vs *New* contrasts, contrasts are reported at a stricter threshold of p < 0.05, *voxel-wise* FWE due to the large clusters obtained at cluster-corrected thresholds. Peak coordinates are reported in MNI coordinates. Hemisphere (H) reported as L/R (Left/Right). ACC: anterior cingulate cortex, MTG: Middle Temporal Gyrus, PCC: posterior cingulate cortex.

| Contrast | Anatomical region | H | Coordinates |  |  | t-value | p-value | Cluster size<br>(voxels) |
| --- | --- | --- | --- | --- | --- | --- | --- | --- |
|  |  |  | x | y | z |  |  |  |
| ***Rel* > *Unrel*** | ACC | L | -6 | 50 | -1 | 5.32 | 0.000 | 343 |
|  | Middle Occipital Gyrus | L | -15 | -100 | 5 | 5.29 | 0.003 | 146 |
|  | Middle Occipital Gyrus | L | -39 | -70 | 35 | 4.96 | 0.000 | 244 |
|  | MTG | L | -51 | -43 | -4 | 4.90 | 0.018 | 95 |
|  | PCC | L | -3 | -46 | 29 | 4.86 | 0.004 | 134 |
|  | Lingual Gyrus | R | 18 | -85 | -10 | 4.79 | 0.002 | 157 |
| ***Unrel* > *Rel*** | Inferior Occipital Gyrus | L | -45 | -70 | -4 | 5.78 | 0.028 | 84 |
| ***Old* > *New*** | Inferior Parietal Lobule | L | -39 | -55 | 44 | 12.59 | 0.000 | 1388 |
|  | Middle Frontal Gyrus | L | -42 | 8 | 53 | 10.69 | 0.000 | 754 |
|  | MTG | L | -60 | -40 | -4 | 7.90 | 0.000 | 41 |
|  | MTG | L | -63 | -25 | -7 | 5.04 | 0.028 | 1 |
|  | Angular Gyrus | R | 36 | -64 | 41 | 6.76 | 0.000 | 141 |
|  | Posterior-Medial Frontal | L | -6 | 20 | 50 | 6.17 | 0.000 | 41 |
| ***New* > *Old*** | Postcentral Gyrus | L | -45 | -25 | 62 | 6.22 | 0.005 | 148 |
|  | Postcentral Gyrus | L | -63 | -22 | 23 | 5.52 | 0.009 | 129 |
|  | Inferior Temporal Gyrus | R | 51 | -61 | -4 | 4.62 | 0.029 | 96 |

New judgments elicited more significant activity than Old judgments (irrespective of whether the pictures were *Related* or *Unrelated* at encoding) in two clusters in the left postcentral gyrus and the right inferior temporal gyrus (**Figure 5A**, **Table 3**). Old judgments elicited more significant activity than New judgments in the left inferior parietal lobule, the left posterior medial frontal cortex, the left middle frontal gyrus, the left MTG, and the right angular gyrus.

**Figure 5.**
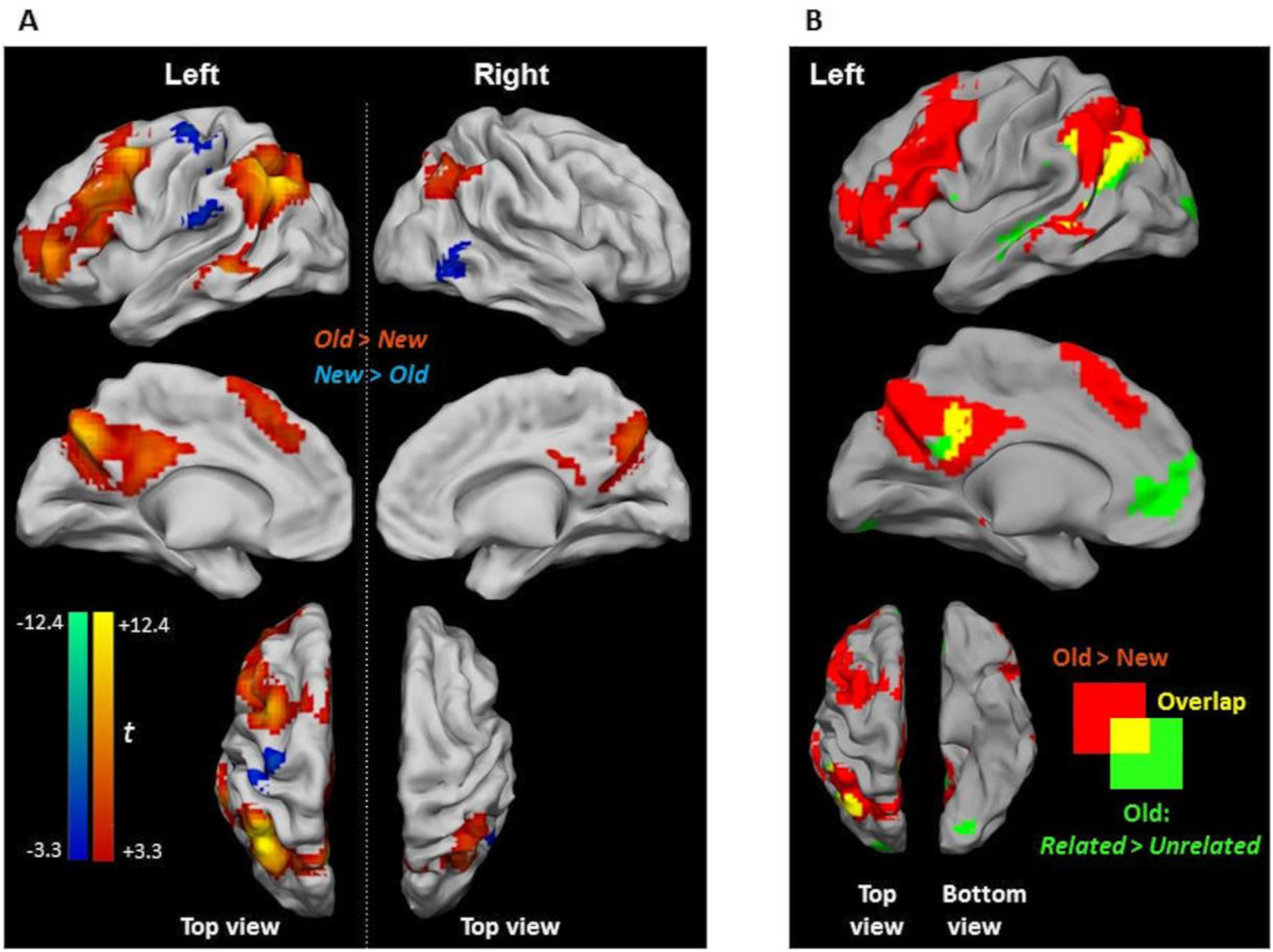
Activity differences during picture-recognition. **A.** Clusters showing significant activity for Old > New (red scale) and for New > Old (blue scale). Due to the large cluster size obtained at cluster-corrected thresholds, contrasts are reported at a stricter threshold of p < 0.05, voxel-wise FWE. **B.** Overlap between Old > New (red) and *Related* > *Unrelated* (green) (from Figure 4C) with the shared area shown in yellow.

A subset of clusters presenting a recognition effect (i.e., Old > New) showed an overlap with the relatedness effect (i.e., Old *Related* > Old *Unrelated*), namely, in the left angular gyrus and PCC (**Figure 5B**). Notably, activity differences in the ACC, which were found in the Old *Related* > Old *Unrelated* contrast due to relatedness, were not observed during recognition (i.e., in the Old > New contrast).

### 3.4 Semantic effects on picture-word recognition

Unlike picture-recognition, each trial of the picture-word recognition task involved the displayed picture along with four words that had to be visually scanned in order to identify the previously seen Old word. The mean accuracy in correctly identifying an Old word was 64.3 ± 15.7%, which was significantly greater than random chance (25%, *U* [61] = 5673, *p* < 0.0001). The mean accuracy to identify words that were part of *Related* picture-word pairs (78.8 ± 15.4%) was significantly higher than for *Unrelated* words (54.4 ± 15.7%, *t* [61] = −12.25, *p* < 0.0001, **Figure 6A**). This effect of relatedness during encoding was also evident in the considerably shorter RT to identify words that were part of *Related* picture-word pairs at encoding (2458 ± 562 ms), in comparison with words from *Unrelated* pairs (3323 ± 640 ms, *t* [60] = 17.85, *p* < 0.0001, **Figure 6B**).

**Figure 6.**
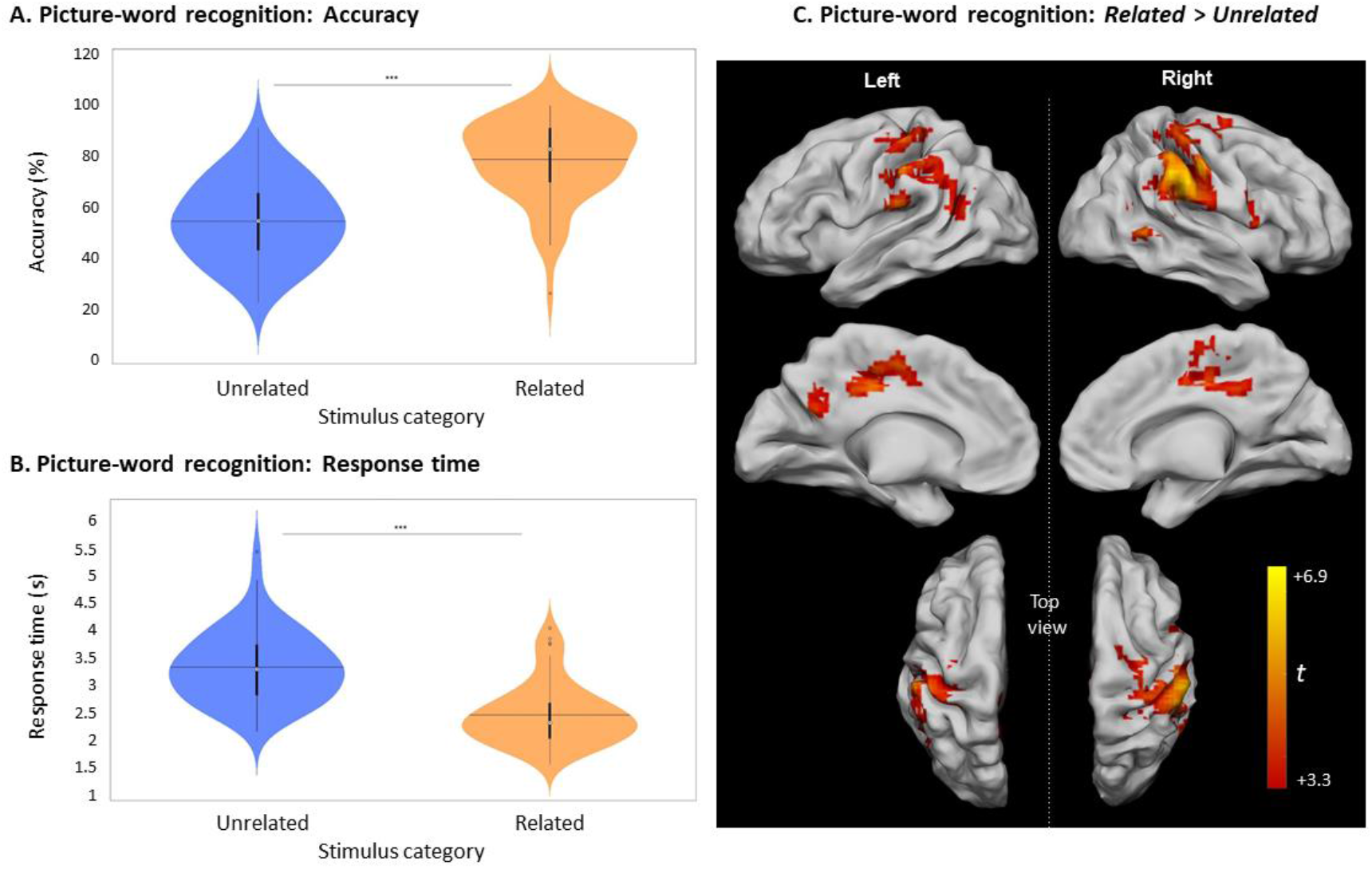
Picture-word recognition according to the judgment about the relatedness of pairs at encoding. **A.** Violin plots of recognition accuracy for picture-word pairs depending on their relatedness during encoding (i.e., *Unrelated* or *Related*) [***: p < 0.001, horizontal black line indicates the mean, thick vertical black bars are a box plot, center white circles represent the median] **B.** Violin plots of Response Times for picture-word pairs depending on their relatedness during encoding [notations as in panel **A**]. **C.** Clusters with significant activity during picture-word recognition for *Related* > *Unrelated*, p < 0.05 FWE-cluster corrected, cluster-forming threshold = 84 voxels (at p < 0.001, uncorr.).

Similar to the encoding and picture-recognition tasks, the neural activity of the last session of picture-word recognition for *Related* pairs was greater than for *Unrelated* pairs (**Figure 6C**, **Table 4**). Activity differences were present in the left inferior parietal lobule, the right supramarginal gyrus, the left middle cingulate cortex (MCC), the right putamen, and the right cerebellum. There was no significant activity difference for the contrast Old *Unrelated* > Old *Related*.

**Table 4.** Picture-word recognition: Peak coordinates for *Related* vs. *Unrelated*. Clusters identified at threshold of p < 0.05, FWE-cluster-corrected. Cluster-forming threshold = 94 voxels (at p < 0.001 uncorr.) Peak coordinates are reported in MNI coordinates. Hemisphere (H) reported as L/R (Left/Right). MCC: middle cingulate cortex

| Contrast | Anatomical region | H | Coordinates |  |  | *t*-value | *p*-value | Cluster size |
| --- | --- | --- | --- | --- | --- | --- | --- | --- |
|  |  |  | x | y | z |  |  |  |
| *Rel > Unrel* | Inferior Parietal Lobule | L | -57 | -25 | 44 | 6.90 | 0.000 | 717 |
|  | Supramarginal Gyrus | R | 60 | -37 | 38 | 6.47 | 0.000 | 1065 |
|  | MCC | L | -3 | -22 | 41 | 5.58 | 0.000 | 584 |
|  | Putamen | R | 33 | 2 | 8 | 4.70 | 0.004 | 158 |
|  | Cerebellum (IV-V) | R | 21 | -43 | -19 | 4.17 | 0.033 | 94 |
| *Unrel > Rel* | Nothing survived after correction |  |  |  |  |  |  |  |

**Figure 7** shows the overlap of regions exhibiting semantically modulated activity during encoding, picture recognition, and picture-word recognition. The semantic modulations during encoding and picture recognition showed an overlap at multiple regions including the left angular gyrus, PCC, and the ACC. However, both encoding and picture recognition showed a limited overlap with picture-word recognition in the anterior aspects of the angular gyrus and the PCC (encoding ⋂ picture-recognition ⋂ picture-word recognition).

**Figure 7.**
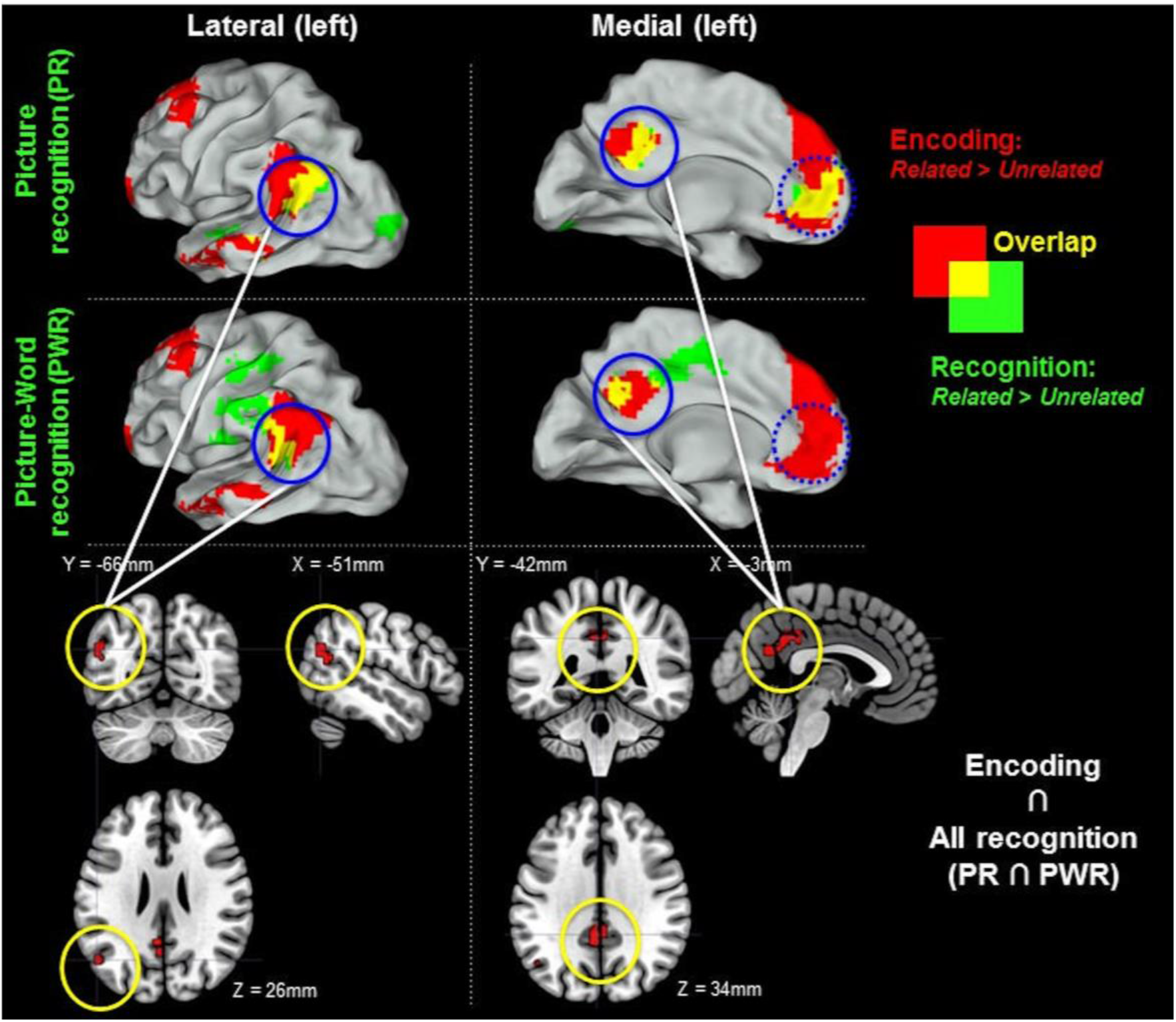
*Related* > *Unrelated* during encoding and recognition. Regions with increased activity for *Related* > *Unrelated* at encoding are shown in red in the upper two rows. Recognition-related activity increases for (Old) *Related* > (Old) *Unrelated* are shown in green: (i) for picture recognition (PR) in the first row and (ii) for picture-word recognition (PWR) in the middle row. The overlap between encoding and recognition-related activity is indicated in yellow. Solid blue circles indicate regions of activity overlaps between encoding and all recognition (encoding ⋂ PR ⋂ PWR). The shared overlaps are shown in sectional views in the bottom row. Dotted blue circles indicate overlaps between encoding and either PR or PWR but not both. There was no overlap in the right hemisphere. The figure shows clusters whose peak survived a size-threshold for p < 0.05 FWE-corrected (cluster-forming threshold at p < 0.001 uncorr.).

### 3.5 Stimulus language and semantic congruency effects

Finally, we assessed whether the choice of stimulus language affects the judgments of semantic congruency during encoding (i.e., *Related* and *Unrelated*) and the associated behavioral and neural consequences for recognition. The findings described thus far were main effects obtained by pooling together all participants despite differences in the chosen stimulus language. The key issue here was whether the subgroups of individuals who chose different languages made equivalent contributions to the shared main effects.

To assess the influence of language choice, we focused on the two largest subgroups consisted of individuals who chose German (GER, n = 42) and who chose French (FR, n = 18) (see Supplementary Table s6 for full details of the sample sizes). As described in **Table 1**, the FR and GER subgroups were similar in gender ratio, age, education, language ability, and cognitive status (MMSE scores). Therefore, possible differences in encoding and recognition processes between these subgroups cannot be merely attributed to differences in the types of individuals that selected each language. For clarity, all analyses described below were limited to individuals with valid fMRI datasets, namely, 41 in the GER subgroup and 17 in the FR subgroup (see Supplementary Table s6).

#### 3.5.1. Limited influence of language on behavioral effects of semantic congruency

During encoding, the picture-word pairs were categorized as being *Related* or *Unrelated* with high agreement to the pre-designed categories by both the FR subgroup (94.04 ± 3.6%) and the GER subgroup (94.51% ± 3.81%) (*U* [57] = 480.5, *p* = 0.72). These matched judgments were a validation that the stimulus sets prepared for the study (Supplementary Tables s1-s5) were semantically equivalent for the FR and GER subgroups.

As described in prior sections, the judgments about the semantic relatedness of the picture-word pairs during encoding (i.e., *Related* or *Unrelated*) were a major determinant of recognition accuracy and RTs during encoding and recognition. As shown in **Figure 8**, the magnitude of these semantic congruency effects varied between tasks, where the effects on accuracy (**Figure 8A**) and RTs (**Figure 8B**) in the picture-word recognition task were substantially larger and more variable than for the encoding and picture-recognition task. The influence of language on these effects of semantic congruency was evaluated with a mixed ANOVA with factors Relatedness [*Related*, *Unrelated*] x Language [FR, GER] as summarized in **Table 5**. Relative to the prominent effects of semantic relatedness (i.e., congruency) on behavior across tasks, language did not have any statistically detectable effects. Furthermore, the size of the semantic congruency effects was comparable between the FR and GER subgroups (**Figure 8**) as confirmed by the non-significant statistical interaction between Relatedness and Language for all the tasks.

**Figure 8.**
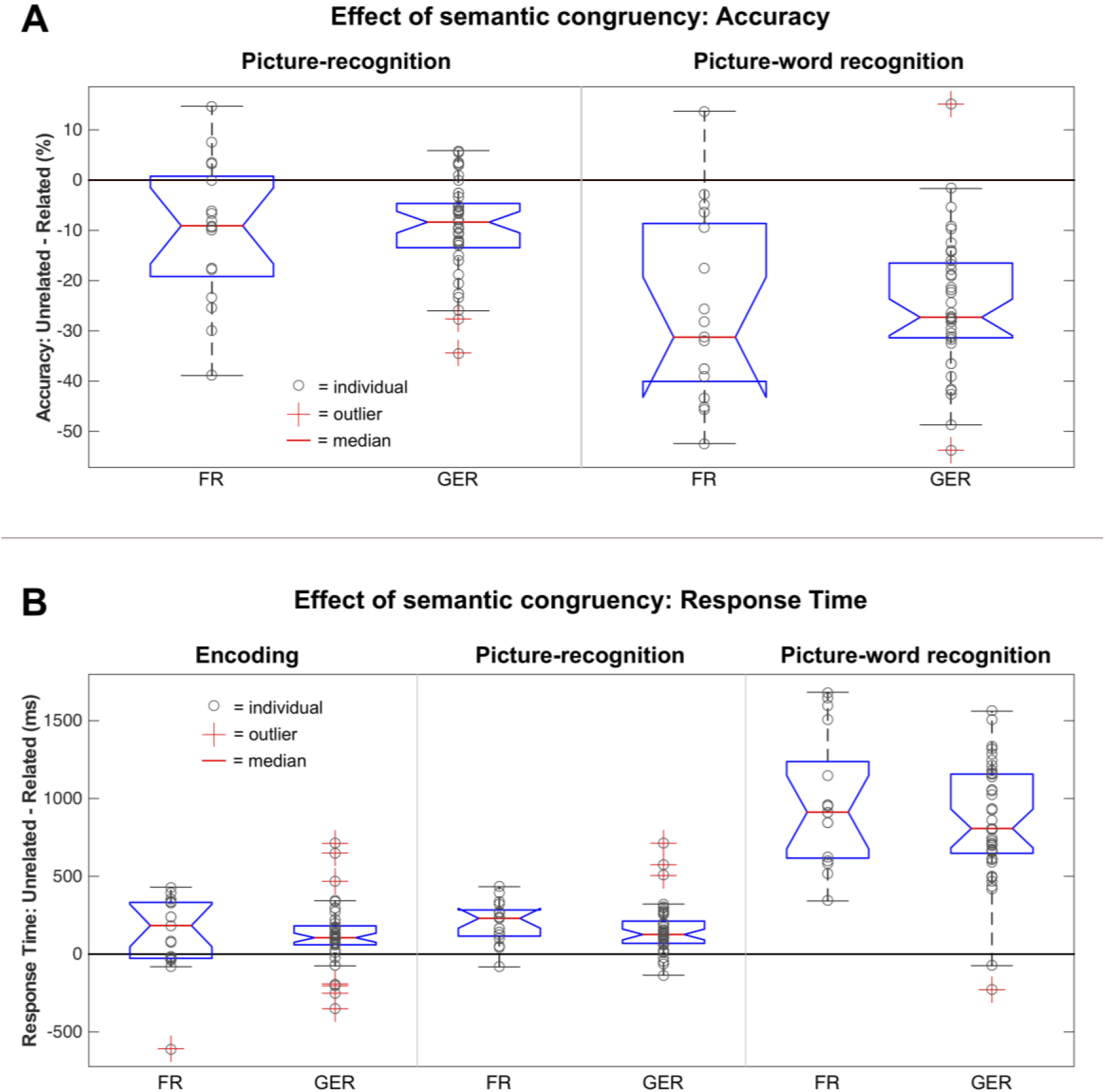
Modulation of semantic congruency effects on behavior by language. **A.** Boxplot of differences in recognition accuracy for *Unrelated* vs *Related* stimuli based on the choice of language (FR = French, GER= German) during picture-recognition (left) and picture-word recognition (right). **B.** Boxplot of differences in Response Times for *Unrelated* vs *Related* stimuli based on the choice of language (FR = French, GER= German) during encoding (left) picture-recognition (middle) and picture-word recognition (right).

**Table 5.** Modulation of semantic congruency effects on behavior by language. Mean accuracy (%) and mean Response Times (ms) for *Unrelated* and *Related* stimuli in the different tasks (Encoding, Picture-recognition, Picture-word recognition) for the French (FR) and German (GER) subgroups (shown as mean ±-standard deviation (SD)). For each task, the behavioral measures were assessed with a mixed ANOVA with factors Relatedness and Language (see text). The corresponding *F* values and *p* are reported on each row (df = degrees of freedom). Statistically significant effects are highlighted in yellow, and non-significant effects in orange.

| *Measure* | *FR*<br>(mean ± SD) | *GER*<br>(mean ± SD) | *Relatedness*<br>df: 1,56 | *Language*<br>df: 1,56 | *Rel.\*Lang.*<br>df: 1,56 |
| --- | --- | --- | --- | --- | --- |
| Encoding |  |  |  |  |  |
| Response time | *Unrel:* 1983 ± 633 ms<br>*Rel:* 1854 ± 726 ms | *Unrel:* 1816 ± 267 ms<br>*Rel:* 1690 ± 263 ms | F = 16.21<br>*p* = 0.002 | F = 1.92<br>*p* = 0.17 | F = 0.002<br>*p* = 0.9661 |
| Recognition: Picture |  |  |  |  |  |
| Accuracy | *Unrel:* 81.0 ± 16 %<br>*Rel:* 91.2 ± 11 % | *Unrel:* 83.1 ± 12 %<br>*Rel:* 92.8 ± 6 % | F = 39.81<br>*p* < 0.0001 | F = 0.44<br>*p* = 0.5095 | F = 0.03<br>*p* = 0.8565 |
| Response time | *Unrel:* 1300 ± 276 ms<br>*Rel:* 1098 ± 206 ms | *Unrel:* 1237 ± 224 ms<br>*Rel:* 1079 ± 142 ms | F = 64.41<br>*p* < 0.0001 | F = 0.54<br>*p* = 0.4649 | F = 0.97<br>*p* = 0.3281 |
| Recognition: Picture-word |  |  |  |  |  |
| Accuracy | *Unrel:* 50.0 ± 14 %<br>*Rel:* 76.1 ± 16 % | *Unrel:* 55.7 ± 16 %<br>*Rel:* 80.7 ± 14 % | F = 141.39<br>*p* < 0.0001 | F = 1.85<br>*p* = 0.1788 | F = 0.08<br>*p* = 0.7784 |
| Response time | *Unrel:* 3589 ± 801 ms<br>*Rel:* 2612 ± 671 ms | *Unrel:* 3237± 549 ms<br>*Rel:* 2388 ± 501 ms | F = 263.88<br>*p* < 0.0001 | F = 3.16<br>*p* = 0.0809 | F = 1.28<br>*p* = 0.2621 |

The matched behavioral effects of semantic congruency on episodic recognition for the FR and GER subgroups support the key individualization assumption of our paradigm.

#### 3.5.2 Limited influence of language on neural effects of semantic congruency

The group-level main effects in previous sections were described by whole-brain activity contrasts. However, a comparison of whole-brain activity of the *Related* > *Unrelated* contrast between the subgroups could be underpowered due to the difference in samples sizes between the FR (n=17) and GER (n=41) subgroups. Therefore, we used a region-of-interest (ROI) approach to evaluate the relative contributions of the FR and GER subgroups to the effects of semantic relatedness on neural activity at selected regions.

The ROIs were selected based on the observation that regions in certain common anatomical zones exhibited elevated activity in the *Related* relative to the *Unrelated* condition both during encoding as well as during recognition (see **Figure 7)**. As summarized in **Figure 9**, we selected clusters in the angular gyrus (AG), posterior cingular cortex (PCC), and middle temporal gyrus (MTG) based on the *Related* > *Unrelated* contrast during the encoding task (**Figure 3B**) and during the picture-recognition task (**Figure 4C**). For this contrast, the shared clusters between encoding and picture-word recognition were present only in the AG and PCC but not MTG (**Figure 7**). The influence of language on the activity at these ROIs was assessed with a mixed ANOVA (Relatedness [*Related*, *Unrelated]* x Language [FR, GER]), as summarized in **Table 6** and **Figure 9**.

**Figure 9.**
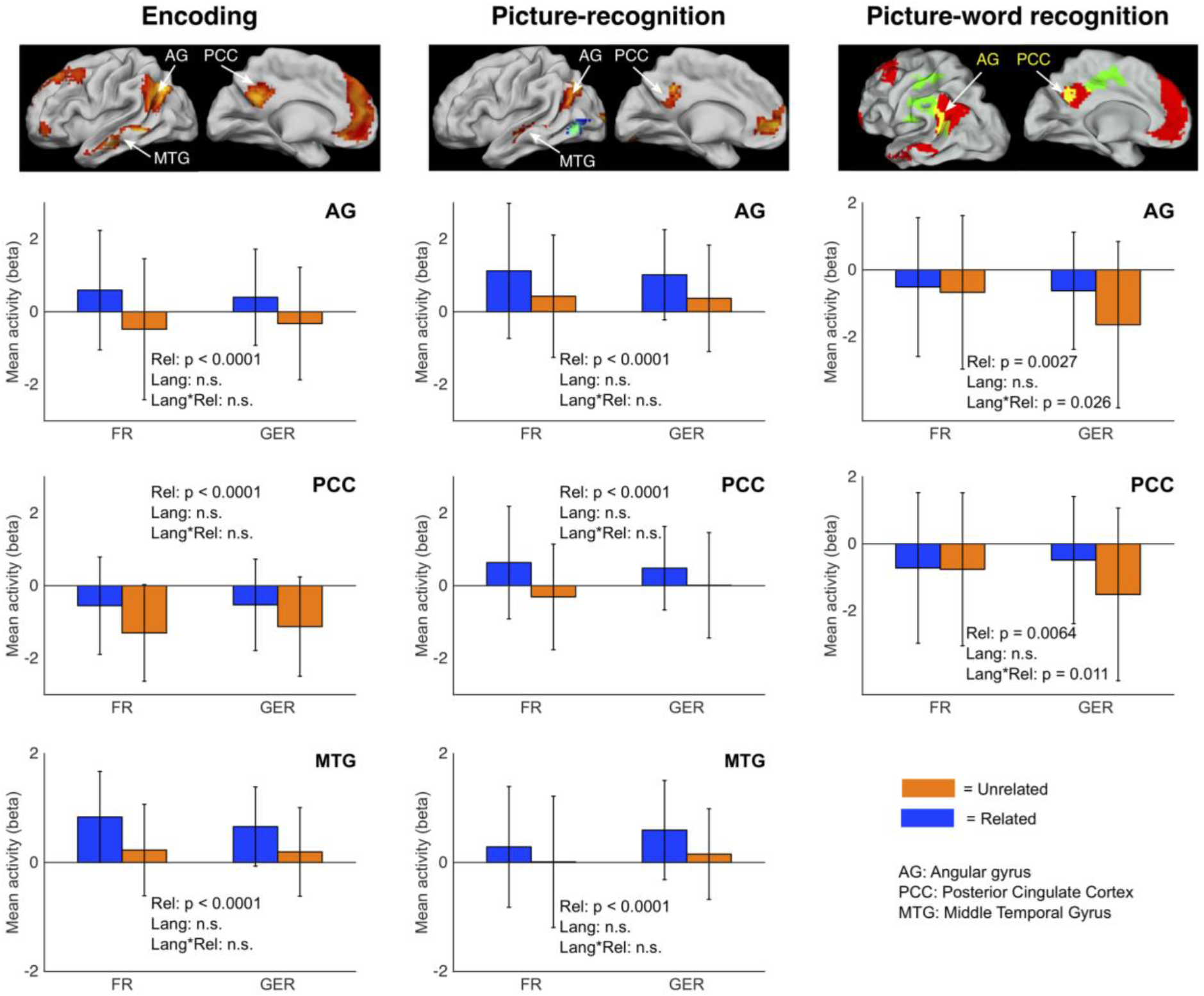
Modulation of semantic congruency effects on neural activity by language. Barplots of mean regional activity during Encoding (left column), Picture-recognition (middle column) and Picture-word recognition (right column) based on Relatedness [*Related, Unrelated*] (abbreviated as Rel) and Language [FR, GER] (abbreviated as Lang). Regions of interest (ROIs) were selected from the group-level *Related* > *Unrelated* contrasts (encoding: Figure 3B; picture-recognition: Figure 4C; picture-word recognition: Figure 7, yellow areas indicating overlap). Error bars indicate the standard deviation. n.s. = not significant (*p* > 0.05, exact p-values are listed in Table 6).

**Table 6.**
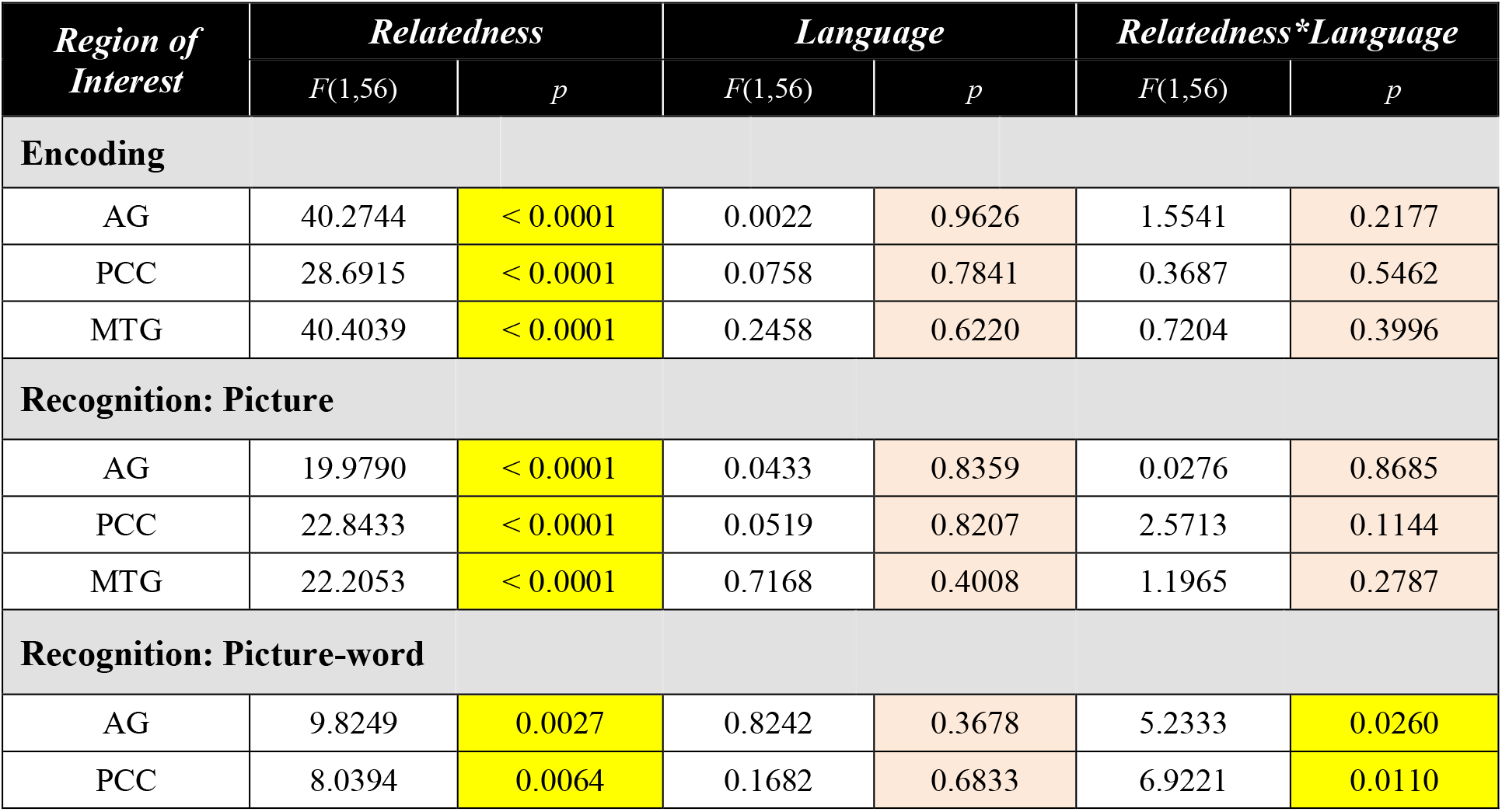
Modulation of semantic congruency effects on neural activity by language. Statistical details of the assessment of regional activity in the different tasks (Encoding, Picture-recognition, Picture-word recognition) with a mixed ANOVA with factors Relatedness and Language (see text). For each region of interest (AG = Angular Gyrus, PCC = Posterior Cingular Cortex, MTG = Middle Temporal Gyrus), the corresponding *F* values and *p* are reported on each row. Statistically significant effects are highlighted in yellow, and non-significant effects in orange.

During encoding (**Figure 9, first column**), semantic congruency had a prominent influence on activity at the AG, PCC and MTG regions of interest but the effect of language was not statistically detectable. Furthermore, the effect of semantic congruency was not detectably modulated by language as confirmed by the non-significant statistical interaction between Relatedness and Language.

In the picture-recognition task (**Figure 9, second column**), the activity differences due to semantic relatedness at the three ROIs were not detectably modulated by language. In the picture-word recognition task (**Figure 9, third column**), language did not have a detectable main effect on activity at the AG and PCC regions but it seemed to modulate the effect of semantic relatedness (indicated by a statistically significant interaction). Specifically, the semantically modulated effects were more prominent for the larger GER subgroup than for the smaller FR subgroup.

Taken together, the behavioral and fMRI findings (albeit with a loss in sensitivity for picture-word recognition) validate the use of stimulus-sets in different languages that are semantically matched in order to elicit comparable semantic congruency effects on encoding and episodic recognition memory across languages.

## 4 Discussion

In the present proof-of-principle study, we sought to validate an individualized paradigm to test the effects of semantic context on episodic object memory where participants could perform the tasks in different participant-chosen languages. This individualization was based on the use of semantically matched stimulus-sets in different languages. With these stimulus-sets, the key assumption was that the effects of categorizing stimuli as being semantically *Related* vs *Unrelated* on encoding and on episodic memory would be effectively independent of the stimulus language.

We here show that with the stimulus-sets used here the paradigm induced a robust effect of semantic context on encoding and recognition behavior. The paradigm also evoked activity in brain regions typically involved in semantic congruency judgments and episodic memory. The semantic congruency effects on behavior and on neural activity during encoding and picture-recognition were comparably present in subpopulations using different task languages (French and German). The behavioral effects of semantic congruency effects were largest for picture-word recognition and were comparably present across languages. However, the large individual variability of the behavioral effects in this latter task reduced the sensitivity in detecting associated neural activity effects at low sample sizes.

Based on these findings, the paradigm suggests a suitability to investigate episodic memory in an inclusive manner in aging, multilingual populations.

### 4.1 Effect of semantic processing on memory encoding

Our paradigm is based on the PWI phenomenon, where the RTs to name a presented object are slower when the distractor word describes a related than an unrelated object (La Heij, 1988, Starreveld and La Heij, 2017). This RT increase is attributed to the increased difficulty in ignoring semantically related information. However, the RT effects in our study were not consistent with such a task difficulty related interpretation.

In the encoding stage of our experiment, the word was not a distractor to be ignored, and participants were instructed to pay attention to the picture and the word. The RTs were shorter for *Related* than for *Unrelated* judgments with these instructions (**Figure 3A**), unlike the PWI effect. If the RT is assumed to be an index of cognitive difficulty (Botvinick et al., 1999, Botvinick et al., 2004), then *Unrelated* judgments should have evoked significantly more neural activity than the *Related* judgments. Specifically, increasing task difficulty has been shown to elicit more brain activity in the parietal, frontal, and cingulate cortices (including ACC, PFC) (Demb et al., 1995, Desai et al., 2006, Gould et al., 2003, Na et al., 2018). However, in our study, neural activity for *Related* items was more significant than for *Unrelated* items across a widespread network of brain areas, including the bilateral angular gyrus, MTG, PCC, ACC, and PFC, thus speaking against task difficulty as the cause for activity changes (**Table 2**, **Figure 3B**).

Two lines of evidence suggest that the activity differences were associated with the effectiveness of memory encoding. During picture-recognition and picture-word recognition, *Related* items were recognized more rapidly and with greater accuracy than items judged to be *Unrelated* (**Figure 4A, B** & **Figure 6A, B**). Also, during picture-recognition and picture-word recognition, *Related* items were associated with relatively higher neural activity relative to the *Unrelated* items over an extended neural network (**Figure 4C**, **Table 3** & **Figure 6C**, **Table 4**). Therefore, relatedness had a significant impact on memory encoding. This interpretation is in line with prior work demonstrating that increased activity during encoding and retrieval of associative memory is associated with better memory performance (Kukolja et al., 2009a, Kukolja et al., 2009b).

### 4.2 Accounting for semantically-mediated recognition

Our findings are consistent with prior studies that have consistently identified the ventral posterior parietal cortex (including the left angular gyrus and MCC/PCC) as having a role in memory processing, even though its functional role remains debated (Davey et al., 2016, Shimamura, 2011).

In light of these prior studies, our findings suggest a straightforward account of how the *Related*/*Unrelated* judgments during encoding might influence memory judgments in our paradigm. When picture-word pairs were judged to be subjectively *Related*, we assume that the participants could successfully identify a *binding* context between a picture and a word. However, when a picture-word pair was judged to be *Unrelated*, we assume that the search for a binding context was unsuccessful. The binding context is internally-generated information that was not contained directly in the stimuli but already present in semantic memory. During picture/picture-word recognition, the binding context facilitates access to encoded item representations. This is akin to the “semantic mediator effectiveness hypothesis” (Carpenter, 2009, Carpenter, 2011, Pyc and Rawson, 2010), where the authors propose facilitation of memory retention when information mediating the association of a cue with a target is present. In our task, this semantically-mediated access might be absent for the *Unrelated* items. Due to these binding-context differences, *Related* items would therefore have a more substantial representation in memory than the *Unrelated* items and should, in turn, lead to faster retrieval and elicit higher neural activity.

Two lines of evidence support the above account. In the picture-word recognition task, the RTs for *Related* items were considerably shorter (~800ms) than for *Unrelated* items. The BC hypothesis suggests an explanation for this substantial behavioral difference. In our task design, picture-word recognition was preceded by the picture-only recognition task, where a picture was presented in isolation. Despite being seen in isolation, when an Old *Related* picture is successfully recognized, the associated binding context might lead to the automatic retrieval of the paired word even though the task does not require it (also see (Carpenter, 2011)). Due to this automatic retrieval, the neural representations of these *Related* words might be more activated than for the *Unrelated* words. This difference in pre-activation might, in turn, affect performance in the subsequent picture-word recognition task. Specifically, it might facilitate more rapid access and recognition of the correct *Related* word from the four presented options than for the *Unrelated* words. In qualitative support of this possibility, during debriefing, 79% of the participants reported that they often remembered the word paired with the picture displayed in the picture (only)-recognition task. This was notable since 69% of the participants also reported that they did not use the *Related*/*Unrelated* judgments during encoding as a conscious strategy to remember the seen picture-word items. This observation is in line with recent work (Amer et al., 2020), which showed that co-occurring congruent, as well as non-congruent material (word and picture paired), are often bound together in memory in older adults.

Finally, the increased engagement of the left angular gyrus and the middle/posterior cingulate cortex (**Figure 7**) for *Related* > *Unrelated* items both during encoding and during recognition (i.e., picture-recognition and picture-word recognition) suggests that these regions might be involved in representing the binding context information (as also suggested by (Shimamura, 2011)).

### 4.3 Semantic processes and language choice

The influence of the language of the word stimuli on semantic processing and memory was a critical consideration of interest in the current study. Although language differences could theoretically lead to systematic cognitive differences (e.g., as postulated by the Sapir-Whorf hypothesis (Koerner, 1992)), we found negligible evidence for this in the behavioral and fMRI data. These findings strongly support the conclusion that individualizing the language materials of the task enables the inclusion of participants with different native languages. This is a crucial strength from a public health viewpoint when studying a multilingual population.

Language choice in the present study could theoretically be associated with education level. In Luxembourg, the language of instruction in primary school for all students is German and French. In secondary school, students chose either a technical path (instructed in German) or a general path (instructed in French). Since Luxembourg did not have a university until 2003 (https://www.en.uni.lu/universite/presentation), students typically attended universities in neighboring countries to obtain an advanced degree (Germany, France, or Belgium, instructed in French). Therefore, a participant’s preferred language might be an indirect proxy for socioeconomic and cultural factors as possible confounds. However, in the present study, neither the number of years of education nor neuropsychological test scores were significantly different between the two language sub-groups (**Table 1**).

An important topic for future research is whether differences in the relative proficiency in the multiple languages spoken by the multilingual participants are a predictor of differences in memory performance in our task and cognitive reserve.

### 4.4 Conclusion

We present a task to study semantic context effects on episodic memory that elicits robust behavioral and neural effects and is largely independent of the language used. These findings support the practical applicability of our paradigm for studying semantic congruency effects on memory in multilingual populations. A central guiding theme for this paradigm was to increase individualization and inclusivity, which are of significant public health relevance in heterogeneous populations. Having adapted and unbiased tools is of high relevance. It is even the basis of the concept of precision public health, where groups of populations are studied as units. A key priority for future research is to assess whether and how the level of multilingual capability modulates age-related changes to the neural systems implementing long-term memory.

## 5 Conflict of Interest

The authors declare that the research was conducted in the absence of any commercial or financial relationships that could be construed as a potential conflict of interest.

## 6 Author Contributions

MP, MV, JS, ND, GF, JK contributed to conception of the study, and MP, SV, OR, MV, JK, to its design. MP collected the data. MP, SV, MV performed the behavioral analysis. MP, SV performed the fMRI analysis, interpreted the data, and wrote the first draft of the manuscript. JK, GF and ND revised the text. JK, GF and LH supervised the study. All authors read, and approved the submitted version.

## 7 Funding

This work was supported by the Luxembourg Institute of Health and the Research Centre Jülich. Additional funding by the Marga- and Walter Boll Foundation to GRF is gratefully acknowledged.

## 8 Acknowledgments

We thank Elke Bechholz, Anita Köth, Nadia Caldarelli for technical support during data acquisition, and the participants for their involvement.

## Notes

### Competing Interest Statement

The authors have declared no competing interest.

### Summary of Updates

We have revised section 3.5 to provide a more detailed analysis of the language-related effects on behavior (3.5.1, Table 5, Figure 8), and brain activity using a ROI-based analysis (3.5.2, Table 6, Figure 9).

## References

Abutalebi, J., Della Rosa, P. A., Green, D. W., Hernandez, M., Scifo, P., Keim, R., Cappa, S. F. & Costa, A. 2012. Bilingualism tunes the anterior cingulate cortex for conflict monitoring. Cereb Cortex, 22, 2076–86.

Aguirre, G. K. 2007. Continuous carry-over designs for fMRI. Neuroimage, 35, 1480–94.

Aguirre, G. K., Mattar, M. G. & Magis-Weinberg, L. 2011. de Bruijn cycles for neural decoding. Neuroimage, 56, 1293–300.

Amer, T., Ngo, K. W. J., Weeks, J. C. & Hasher, L. 2020. Spontaneous Distractor Reactivation With Age: Evidence for Bound Target-Distractor Representations in Memory. Psychol Sci, 31, 1315–1324.

Aminoff, E. M., Kveraga, K. & Bar, M. 2013. The role of the parahippocampal cortex in cognition. Trends Cogn Sci, 17, 379–90.

Andersson, J. L., Hutton, C., Ashburner, J., Turner, R. & Friston, K. 2001. Modeling geometric deformations in EPI time series. Neuroimage, 13, 903–19.

Baldivia, B., Andrade, V. M. & Bueno, O. F. A. 2008. Contribution of education, occupation and cognitively stimulating activities to the formation of cognitive reserve. Dement Neuropsychol, 2, 173–182.

Bar, M. 2004. Visual objects in context. Nat Rev Neurosci, 5, 617–29.

Bialystok, E., Craik, F. I. & Freedman, M. 2007. Bilingualism as a protection against the onset of symptoms of dementia. Neuropsychologia, 45, 459–64.

Bialystok, E. & Craik, F. I. M. 2022. How does bilingualism modify cognitive function? Attention to the mechanism. Psychon Bull Rev.

Botvinick, M., Nystrom, L. E., Fissell, K., Carter, C. S. & Cohen, J. D. 1999. Conflict monitoring versus selection-for-action in anterior cingulate cortex. Nature, 402, 179–81.

Botvinick, M. M., Cohen, J. D. & Carter, C. S. 2004. Conflict monitoring and anterior cingulate cortex: an update. Trends Cogn Sci, 8, 539–46.

Brady, T. F., Konkle, T., Alvarez, G. A. & Oliva, A. 2008. Visual long-term memory has a massive storage capacity for object details. Proc Natl Acad Sci U S A, 105, 14325–9.

Brady, T. F., Konkle, T., Gill, J., Oliva, A. & Alvarez, G. A. 2013. Visual long-term memory has the same limit on fidelity as visual working memory. Psychol Sci, 24, 981–90.

Brodersen, K. H., Ong, C. S., Stephan, K. E., & Buhmann, J. M. 2010. The balanced accuracy and its posterior distribution.. 20th International Conference on Pattern Recognition. IEEE.

Buracas, G. T. & Boynton, G. M. 2002. Efficient design of event-related fMRI experiments using M-sequences. Neuroimage, 16, 801–13.

Canini, M., Della Rosa, P. A., Catricala, E., Strijkers, K., Branzi, F. M., Costa, A. & Abutalebi, J. 2016. Semantic interference and its control: A functional neuroimaging and connectivity study. Hum Brain Mapp, 37, 4179–4196.

Carpenter, S. K. 2009. Cue strength as a moderator of the testing effect: the benefits of elaborative retrieval. J Exp Psychol Learn Mem Cogn, 35, 1563–9.

Carpenter, S. K. 2011. Semantic information activated during retrieval contributes to later retention: Support for the mediator effectiveness hypothesis of the testing effect. J Exp Psychol Learn Mem Cogn, 37, 1547–52.

Chung-Fat-Yim, A., Himel, C. & Bialystok, E. 2019. The impact of bilingualism on executive function in adolescents. Int J Billing, 23, 1278–1290.

Crafa, D., Hawco, C. & Brodeur, M. B. 2017. Heightened Responses of the Parahippocampal and Retrosplenial Cortices during Contextualized Recognition of Congruent Objects. Front Behav Neurosci, 11, 232.

Dash, T., Berroir, P., Joanette, Y. & Ansaldo, A. I. 2019. Alerting, Orienting, and Executive Control: The Effect of Bilingualism and Age on the Subcomponents of Attention. Front Neurol, 10, 1122.

Davey, J., Thompson, H. E., Hallam, G., Karapanagiotidis, T., Murphy, C., De Caso, I., Krieger-Redwood, K., Bernhardt, B. C., Smallwood, J. & Jefferies, E. 2016. Exploring the role of the posterior middle temporal gyrus in semantic cognition: Integration of anterior temporal lobe with executive processes. Neuroimage, 137, 165–177.

Demb, J. B., Desmond, J. E., Wagner, A. D., Vaidya, C. J., Glover, G. H. & Gabrieli, J. D. 1995. Semantic encoding and retrieval in the left inferior prefrontal cortex: a functional MRI study of task difficulty and process specificity. J Neurosci, 15, 5870–8.

Desai, R., Conant, L. L., Waldron, E. & Binder, J. R. 2006. FMRI of past tense processing: the effects of phonological complexity and task difficulty. J Cogn Neurosci, 18, 278–97.

Ehri, L. C. & Ryan, E. B. 1980. Performance of bilinguals in a picture-word interference task. J Psycholinguist Res, 9, 285–302.

Eickhoff, S. B., Stephan, K. E., Mohlberg, H., Grefkes, C., Fink, G. R., Amunts, K. & Zilles, K. 2005. A new SPM toolbox for combining probabilistic cytoarchitectonic maps and functional imaging data. Neuroimage, 25, 1325–35.

Flegal, K. E., Marin-Gutierrez, A., Ragland, J. D. & Ranganath, C. 2014. Brain mechanisms of successful recognition through retrieval of semantic context. J Cogn Neurosci, 26, 1694–704.

Folstein, M. F., Robins, L. N., & Helzer, J. E. 1983. The mini-mental state examination. Archives of general psychiatry, 40, 812–812.

Friesen, D. C., Chung-Fat-Yim, A. & Bialystok, E. 2016. Lexical selection differences between monolingual and bilingual listeners. Brain Lang, 152, 1–13.

Gould, R. L., Brown, R. G., Owen, A. M., Ffytche, D. H. & Howard, R. J. 2003. fMRI BOLD response to increasing task difficulty during successful paired associates learning. Neuroimage, 20, 1006–19.

Koerner, E. F. K. 1992. The Sapir-Whorf Hypothesis: A Preliminary History and a Bibliographical Essay. Journal of Linguistic Anthropology, 2, 173–198.

Konkle, T., Brady, T. F., Alvarez, G. A. & Oliva, A. 2010. Conceptual distinctiveness supports detailed visual long-term memory for real-world objects. J Exp Psychol Gen, 139, 558–78.

Kousaie, S. & Phillips, N. A. 2012. Conflict monitoring and resolution: are two languages better than one? Evidence from reaction time and event-related brain potentials. Brain Res, 1446, 71–90.

Kukolja, J., Goreci, D. Y., Onur, O. A., Riedl, V. & Fink, G. R. 2016. Resting-state fMRI evidence for early episodic memory consolidation: effects of age. Neurobiol Aging, 45, 197–211.

Kukolja, J., Thiel, C. M. & Fink, G. R. 2009a. Cholinergic stimulation enhances neural activity associated with encoding but reduces neural activity associated with retrieval in humans. J Neurosci, 29, 8119–28.

Kukolja, J., Thiel, C. M., Wilms, M., Mirzazade, S. & Fink, G. R. 2009b. Ageing-related changes of neural activity associated with spatial contextual memory. Neurobiol Aging, 30, 630–45.

La Heij, W. 1988. Components of Stroop-like interference in picture naming. Mem Cognit, 16, 400–10.

Li, X., Morgan, P. S., Ashburner, J., Smith, J. & Rorden, C. 2016. The first step for neuroimaging data analysis: DICOM to NIfTI conversion. J Neurosci Methods, 264, 47–56.

Liu, T. T. & Frank, L. R. 2004. Efficiency, power, and entropy in event-related FMRI with multiple trial types. Part I: theory. Neuroimage, 21, 387–400.

Liu, T. T., Frank, L. R., Wong, E. C. & Buxton, R. B. 2001. Detection power, estimation efficiency, and predictability in event-related fMRI. Neuroimage, 13, 759–73.

Lowe, C. J., Cho, I., Goldsmith, S. F. & Morton, J. B. 2021. The Bilingual Advantage in Children’s Executive Functioning Is Not Related to Language Status: A Meta-Analytic Review. Psychol Sci, 32, 1115–1146.

Luk, G. 2008. The anatomy of the bilingual influence on cognition: Levels of functional use and proficiency of language. York University, Toronto, Ontario, Canada: Doctoral dissertation.

Luk, G. & Bialystok, E. 2013. Bilingualism is not a categorical variable: Interaction between language proficiency and usage. J Cogn Psychol (Hove), 25, 605–21.

Na, R., Bi, T., Tjan, B. S., Liu, Z. & Fang, F. 2018. Effect of task difficulty on blood-oxygen-level-dependent signal: A functional magnetic resonance imaging study in a motion discrimination task. PLoS One, 13, e0199440.

Nichols, T., Brett, M., Andersson, J., Wager, T. & Poline, J. B. 2005. Valid conjunction inference with the minimum statistic. Neuroimage, 25, 653–60.

Nichols, T. E., Das, S., Eickhoff, S. B., Evans, A. C., Glatard, T., Hanke, M., Kriegeskorte, N., Milham, M. P., Poldrack, R. A., Poline, J. B., Proal, E., Thirion, B., Van Essen, D. C., White, T. & Yeo, B. T. 2016. Best Practices in Data Analysis and Sharing in Neuroimaging using MRI. bioRxiv.

Packard, P. A., Rodriguez-Fornells, A., Bunzeck, N., Nicolas, B., De Diego-Balaguer, R. & Fuentemilla, L. 2017. Semantic Congruence Accelerates the Onset of the Neural Signals of Successful Memory Encoding. J Neurosci, 37, 291–301.

Perquin, M., Schuller, A. M., Vaillant, M., Diederich, N., Bisdorff, A., Leners, J. C., D’Incau, M., Ludewig, J. L., Hoffmann, D., Ulbricht, D., Thoma, S., Dondelinger, R., Heuschling, P., Couffignal, S., Dartigues, J. F. & Lair, M. L. 2012. The epidemiology of mild cognitive impairment (MCI) and Alzheimer’s disease (AD) in community-living seniors: protocol of the MemoVie cohort study, Luxembourg. BMC Public Health, 12, 519.

Perquin, M., Vaillant, M., Schuller, A. M., Pastore, J., Dartigues, J. F., Lair, M. L., Diederich, N. & Memovie, G. 2013. Lifelong exposure to multilingualism: new evidence to support cognitive reserve hypothesis. PLoS One, 8, e62030.

Power, J. D., Barnes, K. A., Snyder, A. Z., Schlaggar, B. L. & Petersen, S. E. 2012. Spurious but systematic correlations in functional connectivity MRI networks arise from subject motion. Neuroimage, 59, 2142–54.

Pyc, M. A. & Rawson, K. A. 2010. Why testing improves memory: mediator effectiveness hypothesis. Science, 330, 335.

Ratcliff, R. 1993. Methods for dealing with reaction time outliers. Psychol Bull, 114, 510–32.

Rosinski, R. R. 1977. Picture-Word Interference Is Semantically Based. Child Development, 48, 643–647.

Rossion, B. & Pourtois, G. 2004. Revisiting Snodgrass and Vanderwart’s object pictorial set: the role of surface detail in basic-level object recognition. Perception, 33, 217–36.

Shimamura, A. P. 2011. Episodic retrieval and the cortical binding of relational activity. Cogn Affect Behav Neurosci, 11, 277–91.

Starreveld, P. A. & La Heij, W. 2017. Picture-word interference is a Stroop effect: A theoretical analysis and new empirical findings. Psychon Bull Rev, 24, 721–733.

Stroop, J. 1935. Studies of interference in serial verbal reactions. Journal of Experimental Psychology, 18, 643–662.

Wilcox, R. R. & Keselman, H. J. 2003. Modern robust data analysis methods: measures of central tendency. Psychol Methods, 8, 254–74.

Williams, B. D., Pendleton, N. & Chandola, T. 2020. Cognitively stimulating activities and risk of probable dementia or cognitive impairment in the English Longitudinal Study of Ageing. SSM Popul Health, 12, 100656.

Zandt, T. V. 2002. Analysis of response time distributions. Stevens’ handbook of experimental psychology, Third Edition. ISBN 9780471214427, DOI 10.1002/0471214426.

