## Supplementary material for "An individualized fMRI protocol to assess semantic congruency effects on episodic memory in an aging multilingual population": Suppl. mat. Tables s1-s7 (from another version of this preprint: version 2, 2022-05-25)

*Supplementary materials*

**Table s1. Stimulus materials: words used in English**

The Memolingua study protocol used 256 words describing common natural and artificial objects. The colored cells correspond to the words used during encoding, and represent also the position (randomly generated) of the correct answer during picture-word recognition.

| N | Stimulus Category | Congruence | Image Name | Word at Encoding | Words at Picture-word recognition | | | |
|---|---|---|---|---|---|---|---|---|
| | | | | | Word 1 | Word 2 | Word 3 | Word 4 |
| 1 | Old | *Unrelated* | I01 | rabbit | rabbit | rat | squirrel | dog |
| 2 | Old | *Unrelated* | I02 | cardigan | sweatshirt | cardigan | clothes | coat |
| 3 | Old | *Unrelated* | I03 | hen | goose | hen | duck | cock |
| 4 | Old | *Unrelated* | I04 | nail | hammer | drill | nail | screw |
| 5 | Old | *Unrelated* | I05 | arrow | crossbow | arrow | bow | target |
| 6 | Old | *Unrelated* | I06 | spoon | fork | table | plate | spoon |
| 7 | Old | *Unrelated* | I07 | candy | caramel | chocolate | candy | licorice |
| 8 | Old | *Unrelated* | I08 | magician | fairy | magician | witch | stick |
| 9 | Old | *Unrelated* | I09 | fingernail | finger | arm | fingernail | hand |
| 10 | Old | *Unrelated* | I10 | hair | hair | bun | beard | mustache |
| 11 | Old | *Unrelated* | I11 | chalet | chalet | hut | house | building |
| 12 | Old | *Unrelated* | I12 | window | window | shutter | curtain | balcony |
| 13 | Old | *Unrelated* | I13 | coffee | tea | drink | beer | coffee |
| 14 | Old | *Unrelated* | I14 | rake | shovel | garden | pickaxe | rake |
| 15 | Old | *Unrelated* | I15 | rock | rock | pebble | stone | mountain |
| 16 | Old | *Unrelated* | I16 | book | text | word | newspaper | book |
| 17 | Old | *Unrelated* | I17 | tooth | bone | skeleton | tooth | jaw |
| 18 | Old | *Unrelated* | I18 | bread | bread | butter | jam | croissant |
| 19 | Old | *Unrelated* | I19 | star | star | sky | night | moon |
| 20 | Old | *Unrelated* | I20 | swallow | zoo | swallow | cage | bird |
| 21 | Old | *Unrelated* | I21 | walnut | almond | pistachio | walnut | hazelnut |
| 22 | Old | *Unrelated* | I22 | palm | oak | maple | palm | fir |
| 23 | Old | *Unrelated* | I23 | soap | shower | bath | towel | soap |
| 24 | Old | *Unrelated* | I24 | cupboard | dresser | cupboard | shelf | wardrobe |
| 25 | Old | *Unrelated* | I25 | carafe | bucket | carafe | bottle | basin |
| 26 | Old | *Unrelated* | I26 | train | bicycle | motorcycle | airplane | train |
| 27 | Old | *Unrelated* | I27 | gun | sword | saber | carbine | gun |
| 28 | Old | *Unrelated* | I28 | lion | giraffe | lion | tiger | monkey |
| 29 | Old | *Unrelated* | I29 | shop | grocery | bakery | shop | butchery |
| 30 | Old | *Unrelated* | I30 | stairs | elevator | floor | ladder | stairs |
| 31 | Old | *Unrelated* | I31 | cathedral | mosque | temple | cathedral | synagogue |
| 32 | Old | *Unrelated* | I32 | heel | heel | shoe | sock | slipper |
| 33 | Old | *Related* | I33 | water | umbrella | water | rain | mud |
| 34 | Old | *Related* | I34 | wool | glove | ice | snow | wool |
| 35 | Old | *Related* | I35 | key | door | lock | safe | key |
| 36 | Old | *Related* | I36 | suit | shirt | suit | jacket | trousers |
| 37 | Old | *Related* | I37 | desk | eraser | desk | phone | stapler |
| 38 | Old | *Related* | I38 | wine | glass | vine | cellar | wine |
| 39 | Old | *Related* | I39 | goal | match | team | goal | stadium |

| | | | | | | | | |
|---|---|---|---|---|---|---|---|---|
| 40 | Old | *Related* | I40 | sailor | sailor | compass | anchor | lifebelt |
| 41 | Old | *Related* | I41 | mandarin | grapefruit | mandarin | orange | juice |
| 42 | Old | *Related* | I42 | stool | sofa | bed | stool | armchair |
| 43 | Old | *Related* | I43 | nurse | hospital | doctor | bacterium | nurse |
| 44 | Old | *Related* | I44 | postman | postman | letter | stamp | mail |
| 45 | Old | *Related* | I45 | net | pond | aquarium | net | algae |
| 46 | Old | *Related* | I46 | drawing | pencil | drawing | school | color |
| 47 | Old | *Related* | I47 | ring | ring | woman | jewelry | necklace |
| 48 | Old | *Related* | I48 | king | poker | ace | king | heart |
| 49 | Old | *Related* | I49 | crane | crane | facade | tool | scaffolding |
| 50 | Old | *Related* | I50 | soup | soup | gravy | forest | recipe |
| 51 | Old | *Related* | I51 | face | eye | nose | face | cheek |
| 52 | Old | *Related* | I52 | pirate | pirate | hook | treasure | flag |
| 53 | Old | *Related* | I53 | farm | fence | obstacle | saddle | farm |
| 54 | Old | *Related* | I54 | hive | swarm | bee | sting | hive |
| 55 | Old | *Related* | I55 | ashes | firefighter | ashes | fire | alarm |
| 56 | Old | *Related* | I56 | sea | ocean | sea | beach | sand |
| 57 | Old | *Related* | I57 | kitchen | meat | kitchen | chef | restaurant |
| 58 | Old | *Related* | I58 | game | game | six | billiards | pawn |
| 59 | Old | *Related* | I59 | road | sidewalk | crossing | road | car |
| 60 | Old | *Related* | I60 | bouquet | rose | vase | plant | bouquet |
| 61 | Old | *Related* | I61 | orchestra | orchestra | piano | guitar | violin |
| 62 | Old | *Related* | I62 | sugar | dessert | biscuit | candle | sugar |
| 63 | Old | *Related* | I63 | clown | acrobat | makeup | clown | circus |
| 64 | Old | *Related* | I64 | child | girl | boy | child | baby |
| 65 | New | not applicable | I65 | | | | | |
| 66 | New | not applicable | I66 | | | | | |
| 67 | New | not applicable | I67 | | | | | |
| 68 | New | not applicable | I68 | | | | | |
| 69 | New | not applicable | I69 | | | | | |
| 70 | New | not applicable | I70 | | | | | |
| 71 | New | not applicable | I71 | | | | | |
| 72 | New | not applicable | I72 | | | | | |
| 73 | New | not applicable | I73 | | | | | |
| 74 | New | not applicable | I74 | | | | | |
| 75 | New | not applicable | I75 | | | | | |
| 76 | New | not applicable | I76 | | | | | |
| 77 | New | not applicable | I77 | | | | | |
| 78 | New | not applicable | I78 | | | | | |
| 79 | New | not applicable | I79 | | | | | |
| 80 | New | not applicable | I80 | | | | | |
| 81 | New | not applicable | I81 | | | | | |
| 82 | New | not applicable | I82 | | | | | |
| 83 | New | not applicable | I83 | | | | | |
| 84 | New | not applicable | I84 | | | | | |
| 85 | New | not applicable | I85 | | | | | |
| 86 | New | not applicable | I86 | | | | | |

| | | | |
|---|---|---|---|
| 87 | New | not applicable | I87 |
| 88 | New | not applicable | I88 |
| 89 | New | not applicable | I89 |
| 90 | New | not applicable | I90 |
| 91 | New | not applicable | I91 |
| 92 | New | not applicable | I92 |
| 93 | New | not applicable | I93 |
| 94 | New | not applicable | I94 |
| 95 | New | not applicable | I95 |
| 96 | New | not applicable | I96 |

**Table s2. Stimulus materials: words used in French**

The Memolingua study protocol used 256 words describing common natural and artificial objects. The colored cells correspond to the words used during encoding, and represent also the position (randomly generated) of the correct answer during picture-word recognition.

| Image Name | Word at Encoding | Words at Picture-word recognition | | | |
|---|---|---|---|---|---|
| | | Word 1 | Word 2 | Word 3 | Word 4 |
| I01 | lapin | lapin | rat | écureuil | chien |
| I02 | gilet | pull | gilet | habit | manteau |
| I03 | poule | oie | poule | canard | coq |
| I04 | clou | marteau | perceuse | clou | vis |
| I05 | flèche | arbalète | flèche | arc | cible |
| I06 | cuillère | fourchette | table | assiette | cuillère |
| I07 | bonbon | caramel | chocolat | bonbon | réglisse |
| I08 | magicien | fée | magicien | sorcière | baguette |
| I09 | ongle | doigt | bras | ongle | main |
| I10 | cheveux | cheveux | chignon | barbe | moustache |
| I11 | chalet | chalet | cabane | maison | immeuble |
| I12 | fenêtre | fenêtre | volets | rideaux | balcon |
| I13 | café | thé | boisson | bière | café |
| I14 | râteau | pelle | jardin | pioche | râteau |
| I15 | rocher | rocher | caillou | pierre | montagne |
| I16 | livre | texte | mot | journal | livre |
| I17 | dent | os | squelette | dent | mâchoire |
| I18 | pain | pain | beurre | confiture | croissant |
| I19 | étoile | étoile | ciel | nuit | lune |
| I20 | hirondelle | zoo | hirondelle | cage | oiseau |
| I21 | noix | amende | pistache | noix | noisette |
| I22 | palmier | chêne | érable | palmier | sapin |
| I23 | savon | douche | bain | serviette | savon |
| I24 | placard | commode | placard | étagère | armoire |
| I25 | carafe | sceau | carafe | bouteille | bassine |
| I26 | train | vélo | moto | avion | train |
| I27 | pistolet | épée | sabre | carabine | pistolet |

| | | | | | |
|---|---|---|---|---|---|
| I28 | lion | girafe | lion | tigre | singe |
| I29 | magasin | épicerie | boulangerie | magasin | boucherie |
| I30 | escaliers | ascenseur | étage | échelle | escaliers |
| I31 | cathédrale | mosquée | temple | cathédrale | synagogue |
| I32 | talon | talon | chaussures | chaussette | pantoufle |
| I33 | eau | parapluie | eau | pluie | boue |
| I34 | laine | gant | glace | neige | laine |
| I35 | clé | porte | serrure | coffre | clé |
| I36 | costume | chemise | costume | veste | pantalon |
| I37 | bureau | gomme | bureau | téléphone | agrafeuse |
| I38 | vin | verre | vigne | cave | vin |
| I39 | but | match | équipe | but | stade |
| I40 | marin | marin | boussole | ancre | bouée |
| I41 | mandarine | pamplemousse | mandarine | orange | jus |
| I42 | tabouret | canapé | lit | tabouret | fauteuil |
| I43 | infirmière | hôpital | docteur | bactérie | infirmière |
| I44 | facteur | facteur | lettre | timbre | courrier |
| I45 | filet | étang | aquarium | filet | algue |
| I46 | dessin | crayon | dessin | école | couleur |
| I47 | bague | bague | femme | bijou | collier |
| I48 | roi | poker | as | roi | cœur |
| I49 | grue | grue | façade | outil | échafaudage |
| I50 | soupe | soupe | sauce | forêt | recette |
| I51 | visage | œil | nez | visage | joue |
| I52 | pirate | pirate | perroquet | trésor | drapeau |
| I53 | ferme | clôture | obstacle | selle | ferme |
| I54 | ruche | essaim | abeille | piqure | ruche |
| I55 | cendres | pompier | cendres | feu | alarme |
| I56 | mer | océan | mer | plage | sable |
| I57 | cuisine | viande | cuisine | chef | restaurant |
| I58 | jeu | jeu | six | billard | pion |
| I59 | route | trottoir | croisement | route | voiture |
| I60 | bouquet | rose | vase | plante | bouquet |
| I61 | orchestre | orchestre | piano | guitare | violon |
| I62 | sucre | dessert | biscuit | bougie | sucre |
| I63 | clown | acrobate | maquillage | clown | cirque |
| I64 | enfant | fille | garçon | enfant | bébé |

**Table s3. Stimulus materials: words used in German**

The Memolingua study protocol used 256 words describing common natural and artificial objects.
The colored cells correspond to the words used during encoding, and represent also the position (randomly generated) of the correct answer during picture-word recognition.

| Image Name | Word at Encoding | Words at Picture-word recognition | | | |
|---|---|---|---|---|---|
| | | Word 1 | Word 2 | Word 3 | Word 4 |
| I01 | Hase | Hase | Ratte | Eichhörnchen | Hund |

| | | | | | |
|---|---|---|---|---|---|
| I02 | Strickjacke | Pullover | Strickjacke | Kleidung | Mantel |
| I03 | Henne | Gans | Henne | Ente | Hahn |
| I04 | Nagel | Hammer | Bohrer | Nagel | Schraube |
| I05 | Pfeil | Armbrust | Pfeil | Bogen | Zielscheibe |
| I06 | Löffel | Gabel | Tisch | Teller | Löffel |
| I07 | Süßigkeiten | Karamell | Schokolade | Süßigkeiten | Lakritz |
| I08 | Zauberer | Fee | Zauberer | Hexe | Zauberstab |
| I09 | Fingernagel | Finger | Arm | Fingernagel | Hand |
| I10 | Haare | Haare | Dutt | Bart | Schnurrbart |
| I11 | Hütte | Hütte | Bude | Haus | Gebäude |
| I12 | Fenster | Fenster | Rolladen | Vorhang | Balkon |
| I13 | Kaffee | Tee | Getränk | Bier | Kaffee |
| I14 | Rechen | Schaufel | Garten | Spitzhacke | Rechen |
| I15 | Felsen | Felsen | Kiesel | Stein | Berg |
| I16 | Buch | Text | Wort | Zeitung | Buch |
| I17 | Zahn | Knochen | Skelett | Zahn | Kiefer |
| I18 | Brot | Brot | Butter | Marmelade | Croissant |
| I19 | Stern | Stern | Himmel | Nacht | Mond |
| I20 | Schwalbe | Zoo | Schwalbe | Käfig | Vogel |
| I21 | Walnuss | Mandel | Pistazie | Walnuss | Haselnuss |
| I22 | Palme | Eiche | Ahorn | Palme | Tanne |
| I23 | Seife | Dusche | Bad | Handtuch | Seife |
| I24 | Wandschrank | Kommode | Wandschrank | Regal | Kleiderschrank |
| I25 | Karaffe | Eimer | Karaffe | Flasche | Schale |
| I26 | Zug | Fahrrad | Motorrad | Flugzeug | Zug |
| I27 | Pistole | Schwert | Säbel | Jagdgewehr | Pistole |
| I28 | Löwe | Giraffe | Löwe | Tiger | Affe |
| I29 | Laden | Lebensmittellade | Bäcker | Laden | Fleischerei |
| I30 | Treppen | Fahrstuhl | Etage | Leiter | Treppen |
| I31 | Kathedrale | Moschee | Tempel | Kathedrale | Synagoge |
| I32 | Absatz | Absatz | Schuh | Socke | Pantoffel |
| I33 | Wasser | Schirm | Wasser | Regen | Schlamm |
| I34 | Wolle | Handschuh | Eis | Schnee | Wolle |
| I35 | Schlüssel | Tür | Schlüsselloch | Tresor | Schlüssel |
| I36 | Anzug | Hemd | Anzug | Jacke | Hose |
| I37 | Schreibtisch | Radiergummi | Schreibtisch | Telefon | Tacker |
| I38 | Wein | Glas | Rebe | Keller | Wein |
| I39 | Tor | Spiel | Mannschaft | Tor | Stadion |
| I40 | Matrose | Matrose | Kompass | Anker | Rettungsring |
| I41 | Mandarine | Pampelmuse | Mandarine | Orange | Saft |
| I42 | Hocker | Sofa | Bett | Hocker | Sessel |
| I43 | Krankenschwester | Krankenhaus | Arzt | Bakterien | Krankenschwester |
| I44 | Postbote | Postbote | Brief | Briefmarke | Post |
| I45 | Netz | Teich | Aquarium | Netz | Alge |
| I46 | Zeichnung | Stift | Zeichnung | Schule | Farbe |
| I47 | Ring | Ring | Frau | Schmuck | Kette |
| I48 | König | Poker | Ass | König | Herz |

| | | | | | |
|---|---|---|---|---|---|
| I49 | Kran | Kran | Fassade | Werkzeug | Gerüst |
| I50 | Suppe | Suppe | Soße | Wald | Rezept |
| I51 | Gesicht | Auge | Nase | Gesicht | Wange |
| I52 | Pirat | Pirat | Papagei | Schatz | Flagge |
| I53 | Bauernhof | Zaun | Hürde | Sattel | Bauernhof |
| I54 | Bienenstock | Schwarm | Biene | Stich | Bienenstock |
| I55 | Asche | Feuerwehrmann | Asche | Feuer | Alarm |
| I56 | Meer | Ozean | Meer | Strand | Sand |
| I57 | Küche | Fleisch | Küche | Chefkoch | Restaurant |
| I58 | Spiel | Spiel | Sechs | billiard | Spielfigur |
| I59 | Straße | Bürgersteig | Kreuzung | Straße | Auto |
| I60 | Strauß | Rose | Vase | Pflanze | Strauß |
| I61 | Orchester | Orchester | Klavier | Gitarre | Geige |
| I62 | Zucker | Nachtisch | Keks | Kerze | Zucker |
| I63 | Clown | Akrobat | Schminke | Clown | Zirkus |
| I64 | Kind | Mädchen | Junge | Kind | Baby |

**Table s4. Stimulus materials: words prepared in Luxembourgish (but not used)**

The Memolingua study protocol used 256 words describing common natural and artificial objects. The colored cells correspond to the words used during encoding, and represent also the position (randomly generated) of the correct answer during picture-word recognition.

| Image Name | Word at Encoding | Words at Picture-word recognition | | | |
|---|---|---|---|---|---|
| | | Word 1 | Word 2 | Word 3 | Word 4 |
| I01 | Hues | Hues | Raat | Kaweechelchen | Hond |
| I02 | Gilet | Pullover | Gilet | Kleeder | Mantel |
| I03 | Hong | Gäns | Hong | Int | Hunn |
| I04 | Nol | Hummer | Buermaschinn | Nol | Schrauf |
| I05 | Feil | Armbrust | Feil | Bou | Zilscheif |
| I06 | Läffel | Forschett | Desch | Teller | Läffel |
| I07 | Bonbon | Kramel | Schockela | Bonbon | Mokuch |
| I08 | Zauberer | Fee | Zauberer | Hex | Stäbercher |
| I09 | Krall | Fanger | Aarm | Krall | Hand |
| I10 | Hoer | Hoer | Chignon | Baart | Schnauz |
| I11 | Chalet | Chalet | Bud | Haus | Gebai |
| I12 | Fenster | Fenster | Lueden | Rido | Balkon |
| I13 | Kaffi | Tei | Gedrenks | Béier | Kaffi |
| I14 | Reech | Schepp | Gaart | Pioche | Reech |
| I15 | Fiels | Fiels | Wak | Steen | Bierg |
| I16 | Buch | Text | Wuert | Zeitung | Buch |
| I17 | Zant | Knachen | Skelett | Zant | Gebéck |
| I18 | Brout | Brout | Botter | Gebees | Croissant |
| I19 | Star | Star | Himmel | Nuecht | Mound |
| I20 | Schmuebel | Zoo | Schmuebel | Käfeg | Villchen |
| I21 | Noss | Mandel | Pistache | Noss | Hieselnoss |
| I22 | Palm | Eech | Ahorn | Palm | Dänn |

| | | | | | |
|---|---|---|---|---|---|
| I23 | Seef | Dusch | Bued | Duch | Seef |
| I24 | Wandschaaf | Kammoud | Wandschaf | Bicherregal | Kleederschaf |
| I25 | Karaff | Eemer | Karaff | Fläsch | Baséng |
| I26 | Zuch | Velo | Moto | Fliger | Zuch |
| I27 | Pistoul | Schwert | Säbel | Flent | Pistoul |
| I28 | Léiw | Giraff | Léiw | Tiger | Af |
| I29 | Buttek | Epicerie | Bäckerei | Buttek | Metzlerei |
| I30 | Trapenhaus | Lift | Stack | Leeder | Trapenhaus |
| I31 | Kathedral | Moschee | Tempel | Kathedral | Synagog |
| I32 | Tallek | Tallek | Schong | Strëmp | Schlapp |
| I33 | Waasser | Prabbeli | Waasser | Reen | Bulli |
| I34 | Woll | Händsch | Äis | Schnéi | Woll |
| I35 | Schlëssel | Dier | Schlass | Kofferfort | Schlëssel |
| I36 | Kostüm | Hiem | Kostüm | Paltong | Box |
| I37 | Büro | Gummi | Büro | Telefon | Agrafeuse |
| I38 | Wäin | Glas | Drauw | Keller | Wäin |
| I39 | Goal | Partie | Equipe | Goal | Stadion |
| I40 | Matrous | Matrous | Kompass | Anker | Rettungsrank |
| I41 | Mandarin | Pampelmousse | Mandarin | Orange | Jus |
| I42 | Hocker | Couche | Bett | Hocker | Fotell |
| I43 | Krankeschwëster | Spidol | Dokter | Microbe | Krankeschwëster |
| I44 | Bréifdréier | Bréifdréier | Bréiw | Timber | Post |
| I45 | Netz | Weier | Aquarium | Netz | Alg |
| I46 | Zeechnung | Bläistëft | Zeechnung | Schoul | Faarf |
| I47 | Rank | Rank | Fra | Schmuck | Ketten |
| I48 | Kinnek | Pouker | Ass | Kinnek | Häerz |
| I49 | Kran | Kran | Fassad | Handwierksgeschir | Stee |
| I50 | Zopp | Zopp | Zooss | Bësch | Rezept |
| I51 | Gesiicht | Aa | Nues | Gesiicht | Bak |
| I52 | Pirat | Pirat | Papagei | Schatz | Fändel |
| I53 | Bauerenhaff | Zonk | Hindernis | Suedel | Bauerenhaff |
| I54 | Stack | Schwarm | Bei | Stach | Beiestack |
| I55 | Aeschen | Pompjee | Äschen | Feier | Alarm |
| I56 | Mier | Ozean | Mier | Strand | Sand |
| I57 | Kichen | Fleesch | Kichen | Chef | Restaurant |
| I58 | Spill | Spill | Sechs | Billard | Deppchen |
| I59 | Strooss | Greng | Kräizung | Strooss | Auto |
| I60 | Bouquet | Rous | Vas | Planz | Bouquet |
| I61 | Orchester | Orchester | Piano | Gitar | Gei |
| I62 | Zocker | Dessert | Kichelchen | Käerz | Zocker |
| I63 | Clown | Akrobat | Schminke | Clown | Zirkus |
| I64 | Kand | Meedchen | Jong | Kand | Puppelchen |

**Table s5. Pictures selected from normed databases and used in the study (n=96)**

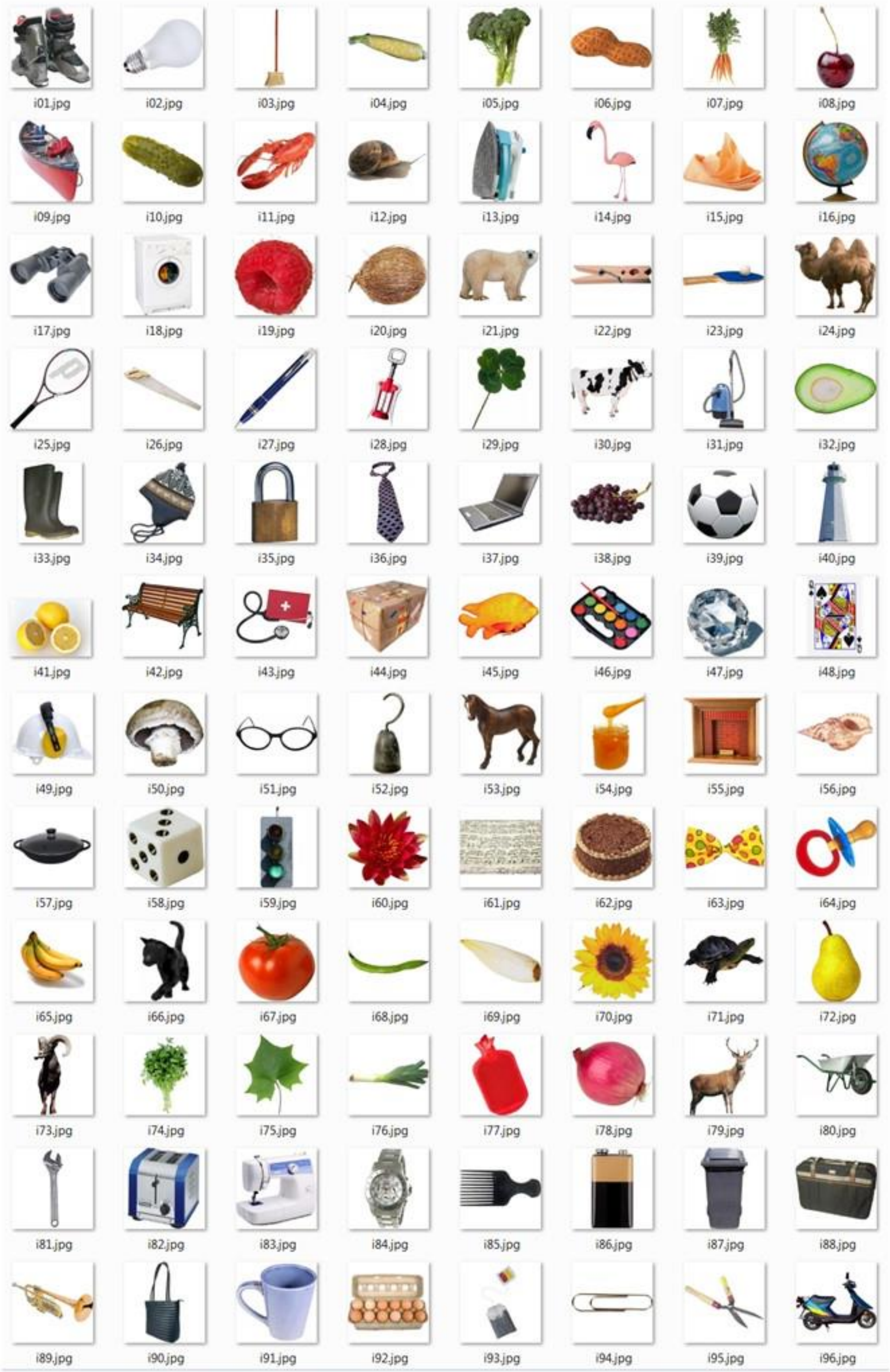

**Table s6. Summary of sample sizes in the different analyses.**

Missing data are indicated by grey cells.

| | Sample sizes | | | | |
|---|---|---|---|---|---|
| **Analysis** | **GER** | **FR** | **ENG** | **LUX** | **Total** |
| **Language choice** | 42 | 18 | 2 | 0 | 62 |
| **Incomplete data** | 1 | 1 | 0 | 0 | 2 |
| **Encoding** | | | | | |
| Accuracy | 42 | 18 | 2 | 0 | 62 |
| RT | 42 | 18 | 2 | 0 | 62 |
| fMRI | 41 | 17 | 2 | 0 | 60 |
| **Picture recogn.** | | | | | |
| Accuracy | 42 | 18 | 2 | 0 | 62 |
| RT | 42 | 18 | 2 | 0 | 62 |
| fMRI | 41 | 17 | 2 | 0 | 60 |
| **Picture-word recogn.** | | | | | |
| Accuracy | 42 | 18 | 2 | 0 | 62 |
| RT | 41 | 17 | 2 | 0 | 60 |
| fMRI | 41 | 17 | 2 | 0 | 60 |
| **Language subgroup comparison** | | | | | |
| Behavior | 42 (41) | 18 (17) | - | - | |
| fMRI | 41 | 17 | - | - | |

**Table s7. Comparison between standard and trimmed mean RTs for the three fMRI sessions.**

***Encoding:***

| Response time: **Trimmed means** | | | Response time: **Means** | | |
|---|---|---|---|---|---|
| Related | Unrelated | statistic | Related | Unrelated | statistic |
| 1749 ± 452 ms | 1877 ± 431 ms *+7.3%* | Z[61]=657.5 p < 0.0001 | 1872 ± 418 ms | 1957 ± 421 ms *+4.5%* | t[61]=3.42 p = 0.0011 Z[61]=494.5 p < 0.0003 |

***Picture recognition:***

| Response time: **Trimmed means** | | | | Response time: **Means** | | | |
|---|---|---|---|---|---|---|---|
| Related Old | Unrel. Old | New | statistic | Related Old | Unrel. Old | New | statistic |
| 1082 ± 158ms | 1247 ± 236ms *+15.2%* | - | Z[61]=905.5 p < 0.0001 | 1157 ± 172ms | 1354 ± 267ms *+17.0%* | - | t[61]=8.86 p < 0.0001 Z[61]=900.5 p < 0001 |
| 1082 ± 158ms | - | 1253.5 ± 211ms *+15.8%* | t[61]=6.17 p < 0.0001 | 1157 ± 172ms | - | 1347 ± 223ms *+16.4%* | t[61]=6.37 p < 0.0001 |
| - | 1247 ± 236ms | 1253.5 ± 211ms *+0.5%* | t[61]=0.20 p = 0.8450 | - | 1354 ± 267ms | 1347 ± 223ms *-0.5%* | t[61]=0.17 p = 0.8684 |

***Picture-Word recognition:***

| Response time: **Trimmed means** | | | Response time: **Means** | | |
|---|---|---|---|---|---|
| Related | Unrelated | statistic | Related | Unrelated | statistic |
| 2458 ± 562 ms | 3323 ± 640 ms *+35.2%* | t[59]=17.85 p < 0.0001 | 2539 ± 517 ms | 3345 ± 598 ms *+31.7%* | t[59]=19.11 p < 0.0001 |

Trimmed means are shown in the left columns and the standard (non-trimmed) means are show in the right columns. For each comparison, the population data were first assessed for normality (using the Kolmogorov-Smirnov test). The *t*-test was used for normal data, and the non-parametric Wilcoxon signed-rank test (*Z* statistic) was used for non-normal data. Note that in two scenarios (Encoding:

*Related* vs *Unrelated*, and Picture-recognition: *Related Old* vs *Unrelated Old*), the trimmed estimates were not normally distributed while the corresponding non-trimmed estimates were normally distributed. Therefore, for comparability, we additionally report the non-parametric statistic (in light gray font) along with the allowed parametric statistic. The values in red show the mean difference in RT. Inter-condition RT differences that were statistically significant with trimmed means remained so with the standard means and differences that were not significant also remained unchanged.
